# TET2 drives 5hmc marking of *GATA6* and epigenetically defines pancreatic ductal adenocarcinoma transcriptional subtypes

**DOI:** 10.1101/2020.10.22.342436

**Authors:** Michael Eyres, Simone Landfredini, Adam Burns, Andrew Blake, Frances Willenbrock, Robert Goldin, Daniel hughes, Sophie Hughes, Asmita Thapa, Dimitris Vavoulis, Aline Hubert, Zenobia D’Costa, Ahmad Sabbagh, Aswin G. Abraham, Christine Blancher, Stephanie Jones, Clare Verrill, Michael Silva, Zahir Soonawalla, Timothy Maughan, Anna Schuh, Somnath Mukherjee, Eric O’Neill

## Abstract

**Background and Aims:** Pancreatic ductal adenocarcinoma (PDAC) is characterised by advanced disease stage at presentation, aggressive disease biology and resistance to therapy resulting in extremely poor five-year survival <10%. PDAC is classified into transcriptional subtypes with distinct survival characteristics, although how these arise is not known. Epigenetic deregulation, rather than genetics, has been proposed to underpin progression but exactly why is unclear and hindered by analysis of clinical samples.

**Methods:** Genome-wide epigenetic mapping of DNA modifications 5-hydroxymethylcytosine (5mc) and 5-hydroxymethylcytosine (5hmc) using oxidative bisulphite sequencing (oxBS). Bioinformatics using iCluster and mutational profiling to identify overlap with transcriptional signatures in FFPE from resected patients and confirmation in vivo.

**Results:** We find that more aggressive squamous-like PDAC subtypes result from epigenetic inactivation of loci including GATA6 that promote differentiated classical-pancreatic subtypes. We show that squamous-like PDAC transcriptional subtypes are associated with greater loss of 5hmc due to reduced expression of the 5mc-hydroxylase TET2. Furthermore, we find that SMAD4 directly supports TET2 levels in the pancreas and classical-pancreatic tumors and loss of SMAD4 expression is associated reduced 5hmc, GATA6 and squamous-like tumors. Importantly, enhancing TET2 stability using Metformin and VitaminC/ascorbic acid (AA) restores 5hmc and GATA6 levels, reverting squamous-like tumor phenotypes and WNT-dependence *in vitro* and *in vivo*.

**Conclusions:** We identify epigenetic deregulation of pancreatic differentiation as an underpinning event behind the emergence of transcriptomic subtypes in PDAC. Our data shows that restoring epigenetic control increases biomarkers of classical-pancreatic tumors and raises the possibility that combination of Vitamin C and Metformin may prolong survival in patients with squamous-like pancreatic cancer.

## INTRODUCTION

There have been several distinct transcriptomic classifications of pancreatic cancer that conform to two main subtypes^1^, and where commonality exists in the recognition of a transcriptomic subtype that performs significantly worse [squamous^2^, basal^3^, Quasi-mesenchymal (QM-PDA) subtype^4^]. Validation and comparison of the three classifications on an independent dataset of 600 patients has confirmed the prognostic differences in the various subgroups^5^. Importantly, the 2-year survival was found to be 23-28% in these “squamous-like” subtypes compared to almost 50 % in the non-squamous subtypes (“pancreatic-classical” subtype). Whereas the prognostic implications of the transcriptomic subtypes have been established, the mechanisms driving the subtypes are much less understood. Genomically, pancreatic cancer is relatively homogeneous, with the main genetic mutations limited to KRAS, CDKN2A, SMAD4 and TP53 and others occurring at low prevalence, but how these genetic events associate with transcriptional signatures is not understood^2^. More recently, there is an increasing recognition of the role of the epigenome in pancreatic carcinogenesis where histone modifications and DNA modifying enzymes, such as DNA-methyltransferases (DNMTs), supporting epigenetic silencing and metastatic dissemination^6, 7^.

Epigenetic silencing results from accumulation of 5mc at specific enhancer or promoter regions (CpG-Islands) that repress gene expression by concentrating the Polycomb Repressive Complexes PRC^8^. Hydroxylation of 5mc to 5hmc allows eventual restoration to cytosine through base excision repair, and is generated by the α-ketoglutarate-dependent enzymes TET1-3^8^. Expression of TET1 is vital for pluripotency in embryonic stem cells and is progressively replaced by TET2 in differentiating endoderm, highlighting marking of distinct gene sets at different stages of development^9^. Recent technological advances have allowed the identification and genomic localisation of 5hmc versus 5mc (e.g. TAPS, TABseq, OxBS^10–12^, however, as a low-frequency mark, this has not been widely possible on clinical material. However, during development 5hmc is specifically enriched at the borders of active enhancer elements and CpG islands and notably devoid of repressive 5mc^13^.

The epigenomic landscape of pancreatic cancer, integrated into a multi-omic approach (RNA-Seq, miRNA-Seq, Exome-Seq, methylation and SNP chips) in patient derived xenografts have demonstrated that classical-pancreatic and basal PDAC subtypes can be identified through epigenomic alterations with the classical subtype associated with genes related to pancreatic differentiation (GATA6, PDX1, BMP2 and SHH)^14–16^. There we present a comprehensive multi-omic analysis of resected tissue for a clinical cohort of pancreatic cancer patients and describes how variations in 5mC and 5hmC actively drive transcriptomic subtypes. Furthermore, we demonstrate how restoration of TET2 activity with metformin and vitamin C (AA) restores 5hmc levels and can lead to switching of the squamous-like transcriptomic subtype to more classical-pancreatic, *in-vitro* and *in-vivo*, opening up avenues for therapeutic intervention.

## RESULTS

### Loss of 5hmc from endoderm differentiation genes in human pancreatic cancer

Squamous PDAC tumors are enriched in KDM6A mutations and 5-methylcytosine (5mc) silencing of GATA6, PDX1 and HNF1B, suggesting epigenetic deregulation is likely to be involved in subtype definition^2^. TET1-3 oxidize 5mc to 5hmc^17^ in a reaction that requires the co-factors α-ketoglutarate (α-KG) and AA in a demethylation cycle^18, 19^. However, 5hmc is also actively maintained in the genome^20–22^ where it suppresses DNMT activity and supports transcription at specific loci^23, 24^. To characterise the (epi)genomic landscape of pancreatic cancer we selected tumor FFPE tissue from treatment-naive PDAC patients undergoing resection at the Churchill hospital Oxford (Supplementary Table 1). We first determined total nuclear 5hmc levels by IHC in tumor and pathological healthy non-affected pancreatic tissue controls (healthy tissue). Staining of healthy tissue (n=3) and histologically normal tissue adjacent to tumors (healthy adjacent n=38) showed moderate to strong positivity throughout the nuclei of cells in β-cells, acini and ducts (Figure 1A). However, 5hmc staining is progressively lost from low to high grade PanINs and PDAC (Figure 1A and Supplementary Figure 1A,B). Next, to determine genome-wide 5mc and 5hmc we adapted the oxBS method^25^ for FFPE and determined differentially methylated and hydroxymethylated probes (DMP/DhMP) on illumina EPIC arrays (Figures 1B and Supplementary Figure 2A-D). Notably, patients that maintained the highest 5hmc had a better disease free survival (Figure 1B). DMP profiles showed the greatest variation in 5mc of pancreatic differentiation genes (e.g. *PROX1* and *PTF1A*) and gene sets for p53, GATA6 and HOXA9 (Supplementary Figure 3A-C), whereas DhMP indicated largest diversity of 5hmc at genes associated with PDAC progression, e.g. MAPK signallling, TGFβ, or genes that acquire 5hmc during pancreatic differentiation^9^ (Supplementary Figure 3D). Genomic localisation of DMP/DhMP revealed that 5hmc is enriched at CpG-islands shores (<2kb adjacent) and inversely correlated with the presence of 5mc within CpG islands (Figure 1D,E)^9, 25^. Notably, protective 5hmc peaks appear to maintain active regulatory regions at genes involved in pancreatic differentiation and β-cell function, NR5A2^26^ and GRB10^27^ (Figure 1F).

**Figure 1:**
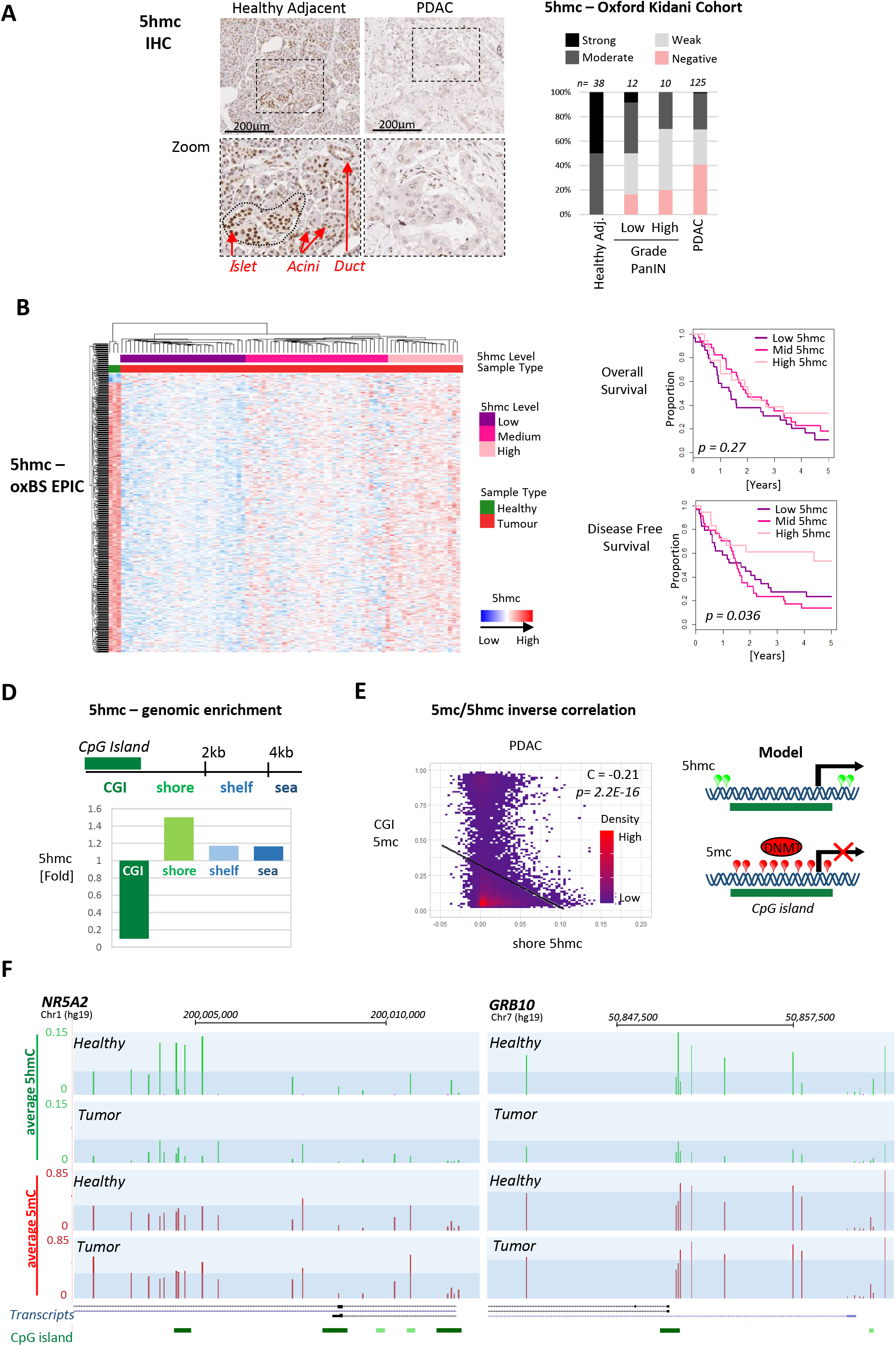
Hydroxymethylcytosine is lost in PDAC. A) IHC staining and quantification of 5hmc in healthy pancreatic tissue, Pancreatic Intraepithelial Neoplasia PanIN lesions and PDAC. B) Hierarchically clustered differentially hydroxymethylated probes (DhMPs) detection on illumina EPIC array. Below, Kaplan-Meier analysis for overall survival and disease free survival with low, moderate and high levels of 5hmc. C) Relative enrichment of 5hmc probes at genomic locations CpG island (CGI), shore, shelf and open sea. D) Correlation of 5mc CGI and 5hmc at CpG shores. Model: 5hmc at the rims of regulatory regions protects from DNMT activity. E) Genomic tracks of *NR5A2* and *GRB10* indicating average 5mc (red) and 5hmc (green) from healthy and tumor.

### Genomic/epigenomic profiling defines two discrete sub-types of pancreatic cancer

To determine the impact of 5hmc we next integrated the epigenetic landscape with genomic mutations determined by focused sequencing for a panel of 52 commonly mutated genes in cancers (Supplementary Figure 4A). The most variable 5mc and 5hmc probes (top 2000) were combined with binary mutation data to cluster multiple data types using iCluster^28^. Bayesian information criteria (BIC) modelling showed that most of the variation between samples could be explained with two equally sized clusters, distinct in their genetic, epigenetic and clinical properties (Figure 2A). Cluster 1 was characterized by low 5hmc, high 5mc and increased incidence of mutations in TP53, SMAD4, RB1 and VGFR2. Patients in Cluster 1 associated with increased rates of recurrence, indicating a more aggressive set of tumors, and poorer overall and disease free survival compared to Cluster 2 (HR=0.53 and 0.50 respectively) (Figure 2B). An mRNA array from matched FFPE tissue scrolls revealed that Cluster 1 displayed increased transcription of key pathways associated with PDAC tumor progression, EMT, angiogenesis, tumor metastasis and cisplatin resistance (Figure 3A). Gene sets enriched in Cluster 1, such as ΔNp63, Myc and Wnt^2^, have also been linked to the squamous-like subtypes, such while Cluster 2 was enriched for gene sets related to terminal pancreatic differentiation^29^ and less aggressive classical-pancreatic subtypes (Figure 3A).

**Figure 2:**
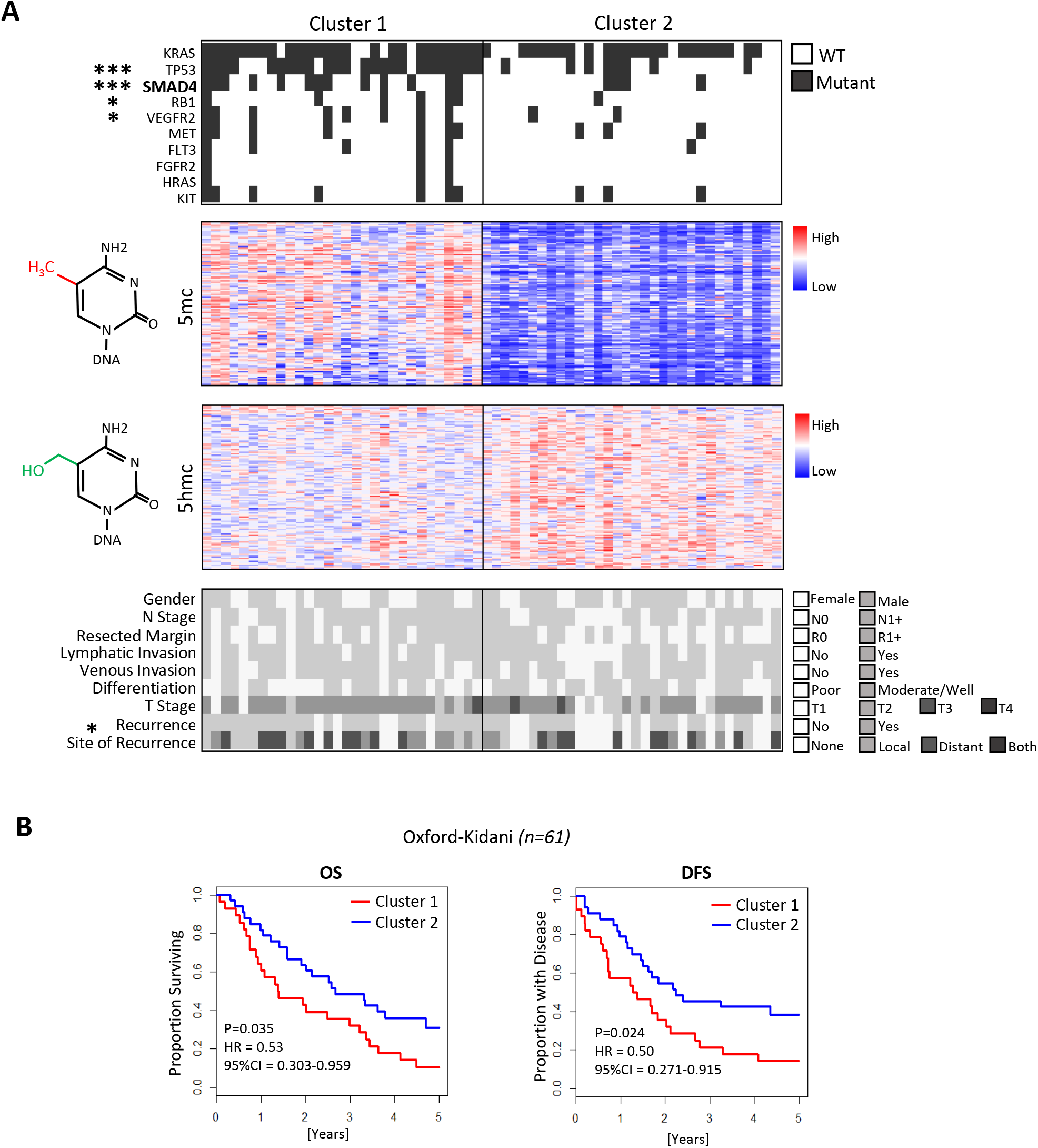
Genetic and Epigenetic clustering identifies two distinct subgroups. A) Integrated (epi)genomic analysis using mutation (Supplementary Figure 4), 5mc and 5hmc data in the Oxford-Kidani Cohort. B) Kaplan-Meier analysis of overall and disease free survival for each cluster.

**Figure 3:**
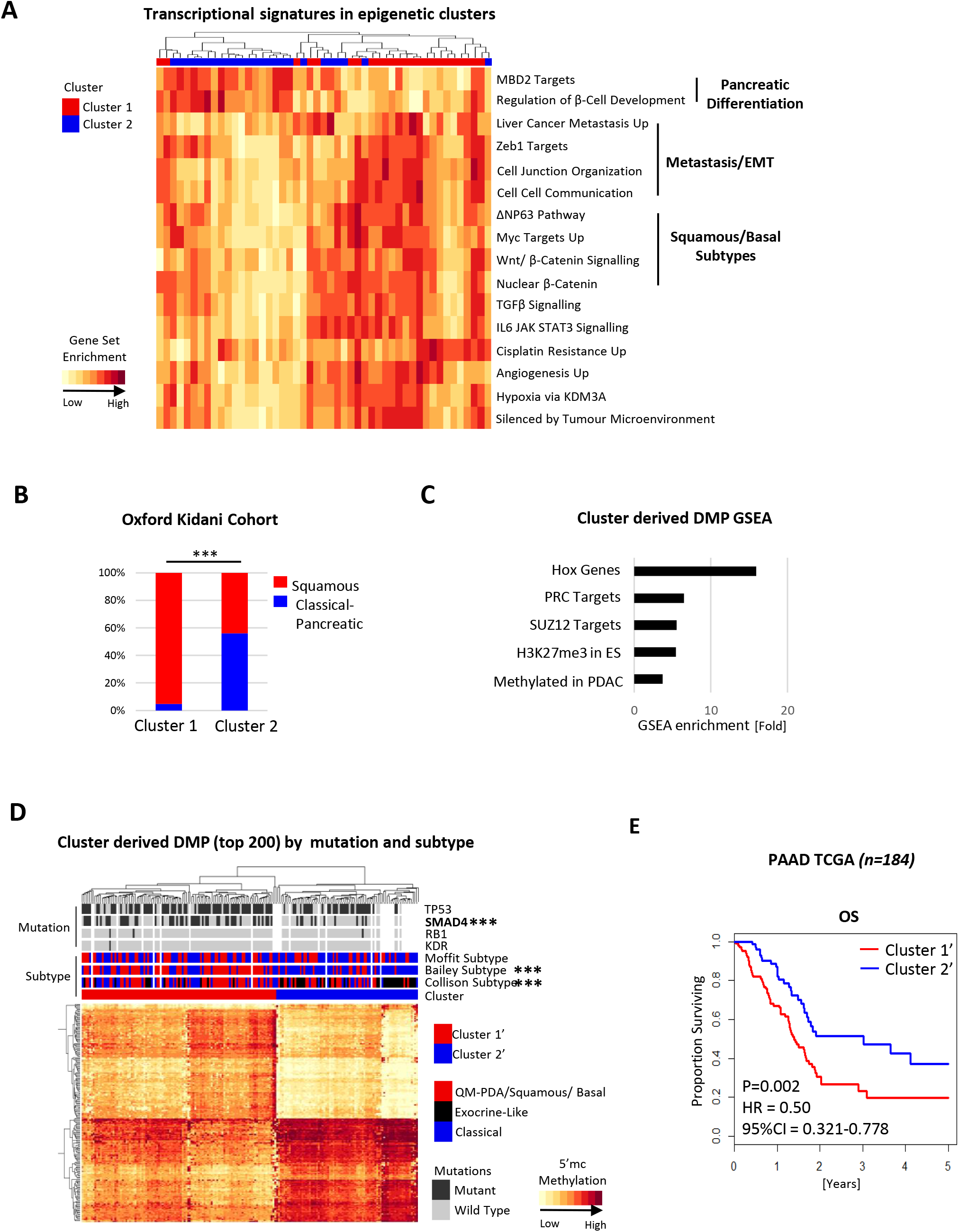
Loss of 5hmc correlates with Squamous subtypes and SMAD4 mutations. A) Differential GSEA between (epi)genetic clusters. B) Distribution of squamous and classical-pancreatic transcriptional subtypes in (epi)genetic cluster defined mRNA signatures. C) GSEA of DMP between (epi)genetic clusters. D) Application of top 200 DMPs between clusters to the TCGA_PAAD cohort. Clusterl’ and 2’ represent the methylome of Clusterl and Cluster2 patients. E) Kaplan-Meier analysis between Cluster1’ and Cluster2’ in the TCGA_PAAD cohort.

To determine overlap with previously described pancreatic subtypes and the currently described (epi)genomic clusters, tumors were classified according to the original PDAC gene sets using mRNA expression data. A gene set was created for each subtype and relative enrichment determined using GSVA across the Oxford-Kidani cohort. Tumors were then hierarchically clustered and assigned subtypes according to the most enriched gene set in each cluster, revealing extensive overlap between aggressive subtypes (Supplementary Figure 4B)^1^. Next, tumors classified as squamous, basal or QuasiMesemchymal-PDA^1^ were grouped (Merged) into a single subtype termed squamous, and remaining subtypes grouped as classical-pancreatic (Supplementary Figure 4B). Interestingly, >95% Cluster 1 tumors classified as a squamous subtype, therefore reflecting the genetic and epigenetic basis of transcriptionally subtyped tumors (Figure 3B). These results were then validated using the 5mc component of the iCluster against available 5mc data in TCGA_PAAD (n=184). DMP analysis indicated that Cluster 1 ‘ tumors had an embryonic stem cells-like epigenome with increased methylation of Hox genes, frequently silenced in pancreatic cancer^30, 31^ and regulated by TET2 during development^32^ (Figure 3C). The top 200 DMPs (5mc methylome) were sufficient to identify Cluster 1’ with squamous-like subtypes in the PAAD_TCGA dataset, enrichment for SMAD4 mutation and reduced overall survival compared to Cluster 2’ patients (HR = 0.50) (Figure 3D, E). Therefore, the epigenetic signature derived from multi-omic clustering of patients is a robust predictor of patient survival and molecular identity.

### TET2 and 5hmc levels are directly correlate with GATA6 expression

TET1 supports pluripotency of embryonic stem cells but becomes gradually depleted and replaced by TET2 as cells differentiate through endodermal intermediates into pancreatic epithelial cells. Concomitantly, 5hmc is high in ES cells and lost as TET1 decreases in progenitors but accumulates again at active regulatory regions of key pancreatic differentiation genes in line with TET2 expression^9^. The endodermal marker GATA6 is critical for pancreatic specification during development, directly activating the Pancreatic and Duodenal homeoboX transcription factor, PDX1. Moreover, GATA6 maintains the classical-pancreatic subtype^33^ and loss of expression is a robust surrogate biomarker for squamous tumors^34, 35^. To identify the route of the epigenetic imbalance between subtypes we next classified 44 PDAC cell lines available from The Cancer Cell Line Encyclopedia (CCLE)^36^ for enrichment of squamous (squamous/basa/QM-PDA) vs classical-progenitor (progenitor/classical) subtypes using GSVA and found lower expression of GATA6 in squamous cell lines correlates with the pancreatic 5hmc hydroxylase TET2 (Supplementary Figure 5A). Supporting this, TET2, but not TET1 or TET3, directly correlates with expression of GATA6 across the Oxford-Kidani Cohort (C = 0.53; p = 8.8e-08, Figure 4A and Supplementary Figure 5B). In line with TET2 expression, classical-pancreatic tumors also display the highest total 5hmc levels suggesting that TET2 and 5hmc are related to GATA6 expression and subtypes (Figure 4B). Strikingly, a single probe in the 3’ of *GATA6* (cg21040686) represents a significant DhMP between squamous and classical-pancreatic tumors (Figure 4C), and which inversely correlates with 5mc of the CGI in *GATA6* and positively correlates with transcript levels across the cohort (Figure 4D).

**Figure 4:**
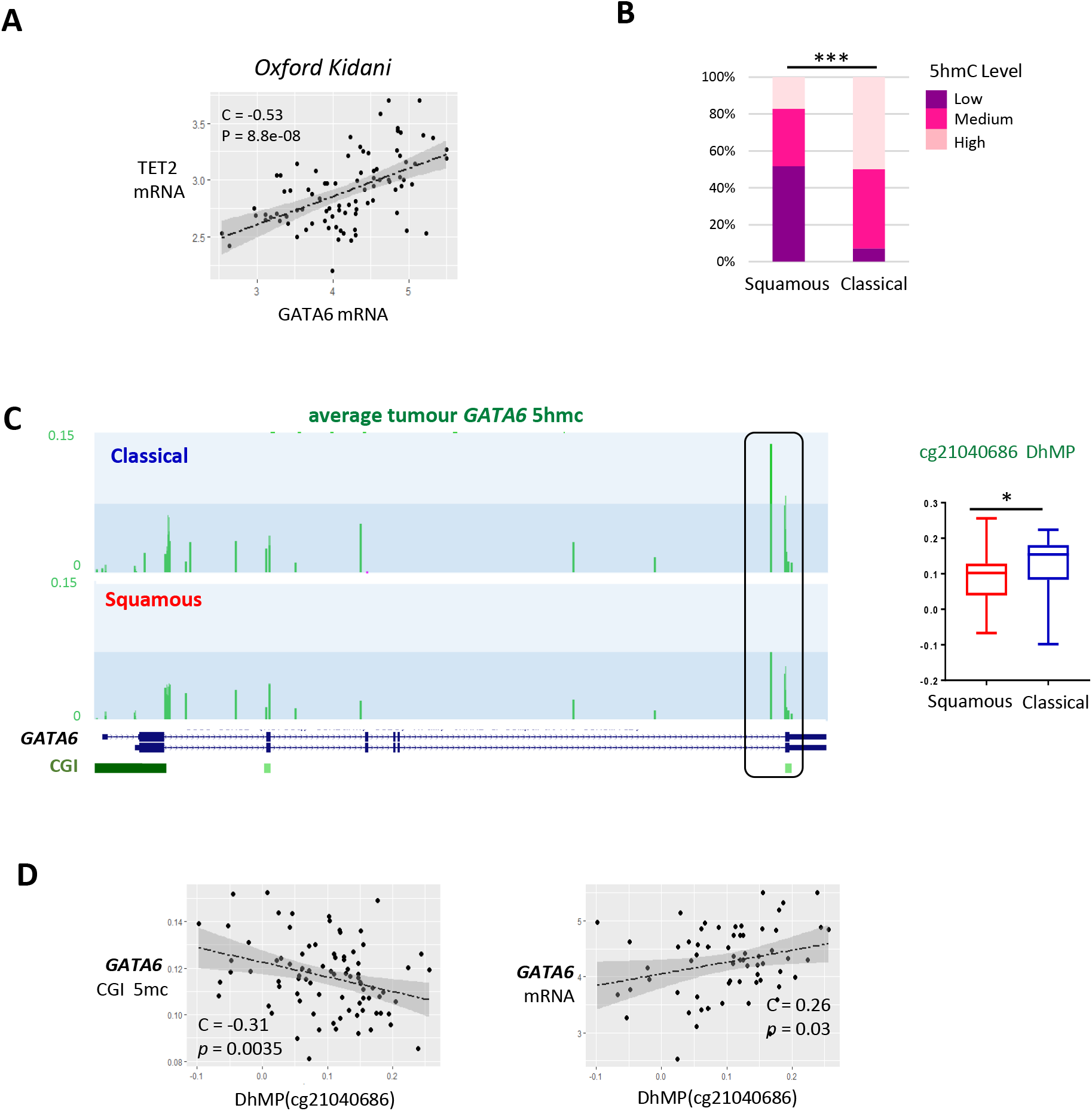
TET2 regulates the Classical-Pancreatic marker GATA6. A) Correlation of *TET2* and *GATA6* expression in the Oxford-Kidani Cohort. B) Distribution of patients with low, moderate and high 5hmc across tumors specific DhMPs in squamous and classical-pancreatic tumors. C) Top: genome tracks showing average 5hmc levels across *GATA6* in classical-pancreatic and squamous tumors. Right, quantification of a single DhMC probe in *GATA6* (cg21040686) with tumor subtypes. D) Left correlation of the DhMP cg21040686 with *GATA6* CGI 5mc and right, with *GATA6* mRNA expression.

### SMAD4 directly regulates TET2, 5hmc and classical pancreatic markers GATA6

Low SMAD4 expression is associated with squamous tumors^4^ (Figure 5A), and we find a highly significant positive correlation between SMAD4 and GATA6 mRNA (C=0.641; *p=* 2.00e-11, Figure 5B). As SMAD4 mutations are associated with Cluster 1 in both Oxford-Kidani and TCGA cohorts (Figures 2A and 3F), we reasoned that loss of SMAD4 may contribute to the initial loss of TET2 and 5hmc, especially as patients with SMAD4 mutations had lower levels of total tumor 5hmc and specifically cg21040686 5hmc (Supplementary Figure 5C). Furthermore, restoration of SMAD4 in tumor cells derived from KSIC (*Kras, Smad4, Ink4A, Cre*)^37^ mice promoted enrichment of transcriptional changes consistent with a less squamous subtype (Figure 5C). ChIPseq data (GSE27526) indicates that TGF-β induces SMAD4 binding to a potential enhancer proximal to the distal 3’ region of *TET2* (DNase hypersensitive site, H3K4me1^+ve^, H3K27ac^+ve^)^38^(Figure 5D). Moreover, we find *SMAD4* expression correlates with *TET2* mRNA across both the Oxford-Kidani (discovery) and TCGA (validation) cohorts, suggesting *TET2* is potentially a SMAD4 target gene (Figure 5E, Supplementary Figure 5D). To test this, we suppressed SMAD4 by siRNA in immortalized ductal epithelial cells (DEC-hTERT) and found this results in decreased TET2 (mRNA and protein), lower global levels of 5hmc, reduction of GATA6 and the marker of pancreatic differentiation PDX1(Figure 5F). Conversely, FLAG-SMAD4 overexpression in PSN1 cells (*SMAD4* deleted)^39^ resulted in reexpression of TET2, increased 5hmc levels and re-establishment of GATA6 and PDX1 (Figure 5F). To determine if TET2 and 5hmc are directly responsible for maintaining the classical-pancreatic transcriptional subtype, we restored TET2 expression in both PANC1 (SMAD4 *wild-type*) and PSN1 squamous cell lines (Figure 5G). In both cases TET2 expression increased 5hmc and raised levels of GATA6, PDX1 and epithelial cadherin (E-cadherin), consistent with a more classical-pancreatic subtype. Thus, while SMAD4 mRNA levels correlate with TET2 expression and GATA6, wild type SMAD4 alone is insufficient to maintain classical-pancreatic subtype, suggesting that additional factors contribute to TET2 levels and activity (e.g. αKG, AA, hyperglycemia and hypoxia).^18, 19, 40, 41^

**Figure 5:**
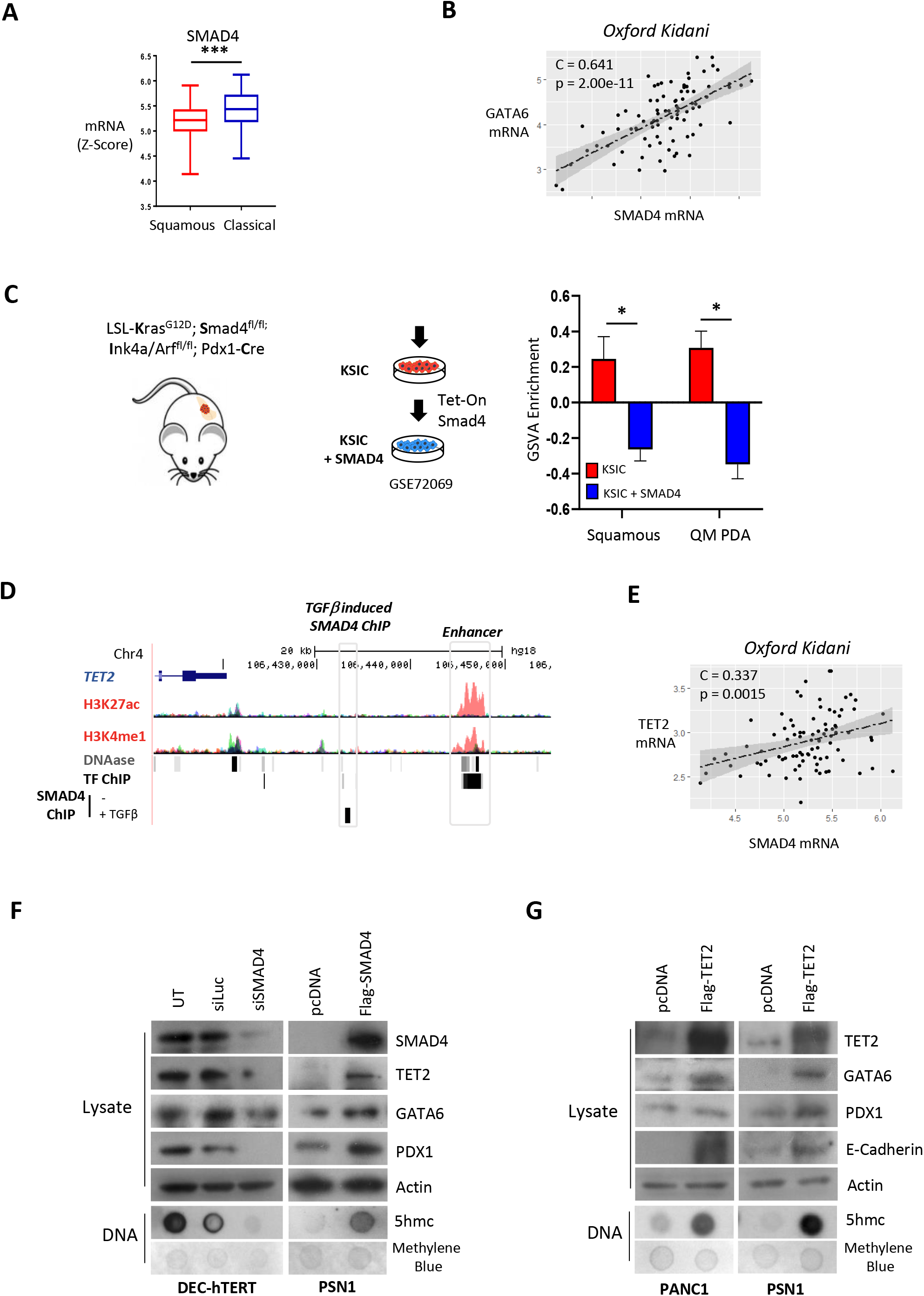
SMAD4 regulates TET2 and GATA6. A) comparative *SMAD4* mRNA expression in tumor subtypes and (B) correlation with *GATA6* expression in the Oxford-Kidani cohort. C) Enrichment of squamous and QM PDA gene sets from RNAseq of tumor cells from LSL-Kras^G12D^; Smad4^fl/fl^; Ink4a/Arf^fl/fl^; Pdx1-Cre transfected with Flag-SMAD4 or control plasmid^27^. D) ChIP-seq dataset (GSE27526) indicating SMAD4 and enhancer binding peaks in the TET2 genomic location following TGFβ stimulation. E) Correlation between *TET2* and *SMAD4* mRNA in Oxford-Kidani and TCGA_PAAD datasets. F) Western blotting of lysates with indicated antibodies of; left, immortalised human Ductal Epithelial Cells-hTERT cells (DEC-hTERT) and transfected with siSMAD4 or non-targeting control; right, PSN1 cells overexpressing Flag-SMAD4 plasmid or control. Global 5hmc was determined by dot blot. MB = methylene blue control. G) Lysates of PSN1 and PANC1 squamous cells overexpressing Flag-TET2 or control and western blotted with indicated antibodies and global 5hmc determined by dot blot.

### Metformin and Ascorbic Acid restore 5hmc and re-expression of GATA6 in squamous-like PDAC

TET2 protein stability is sensitive to reduced AMPK activity associated with hyperglycemia^40^. As the anti-diabetic drug Metformin restores AMPK activity and stabilizes of TET2, we reasoned that treatment with Metformin may also stabilise TET2 in squamous tumors. In PSN1 cells, Metformin (500μM) efficiently stabilises TET2, restores 5hmc and re-establishes GATA6 expression in an AMPK dependent manner, however, physiological relevant doses equivalent to circulating concentration of 20 μm did not (Figure 6A)^42^. Similarly, ascorbic acid (AA) supports TET enzymatic activity by providing reducing power in the hydroxylation reaction^43^. PSN1 cells treated with 500 μm AA displayed a robust three-fold increase in global 5hmc and increased GATA6 expression, whereas lower physiological levels (100 μM)^43^ did not elicit a response (Figure 6B). However, combining low dose AA with the 20 μM Metformin produced a synergistic increase in TET2 and 5hmc levels, leading to re-expression of GATA6 in PSN1, PANC1 and MiaPaca2 squamous cell lines (Figure 6C and Supplementary Figure 6B,C), but did not affect levels in classical-pancreatic or ductal cells that express GATA6 (Supplementary Figure 6D).

**Figure 6:**
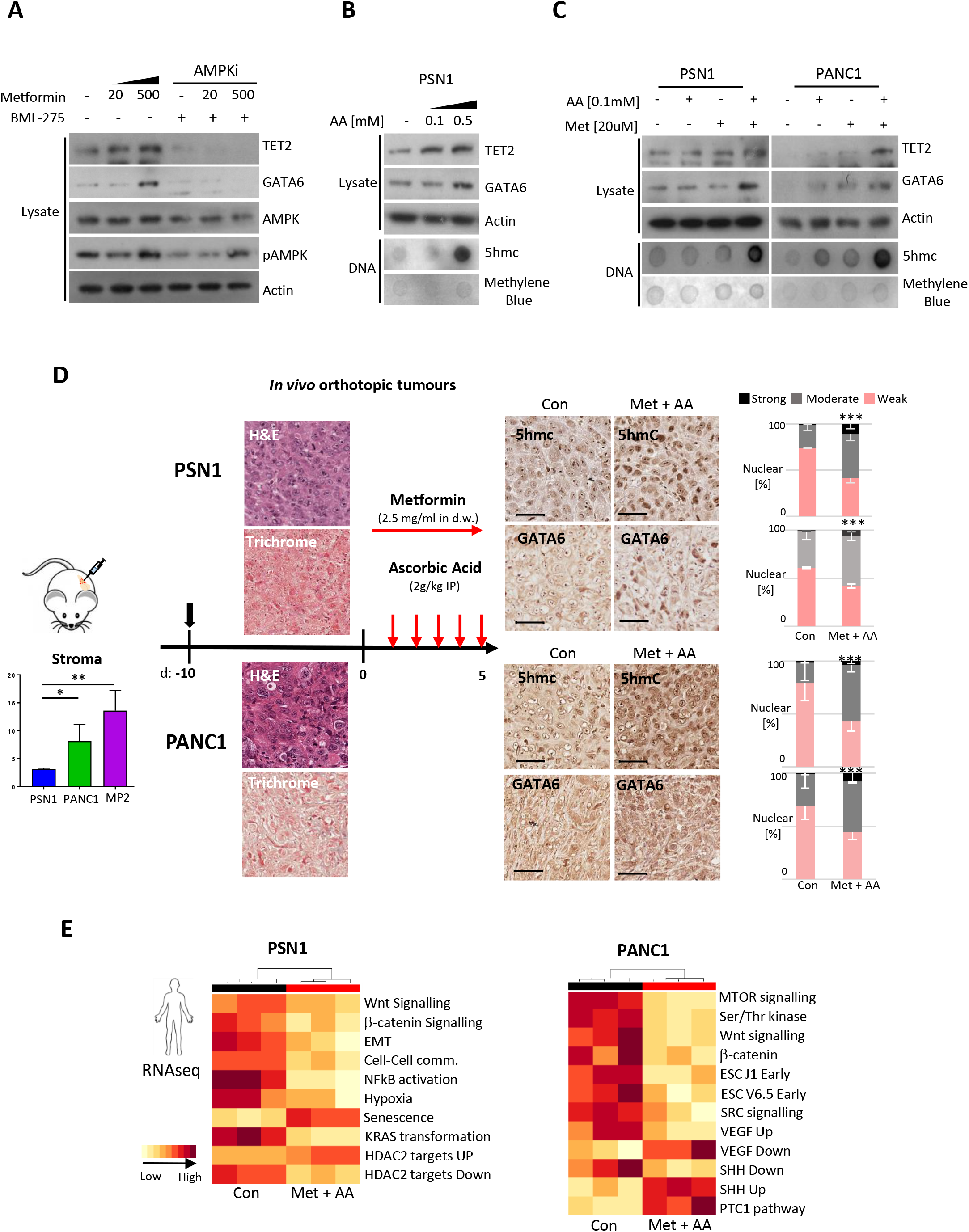
Restoration of 5hmc using Metformin and Ascorbic Acid Alters Subtype Identity. A) Western blots of PSN1 cell lysates after treatment with indicated concentrations of Metformin in the presence or absence of AMPK inhibition. B) Western blots and dot blots of PSN1 cell lysates after treatment with indicated concentration of Ascorbic acid (AA). C) Treatment of PSN1 and PANC1 cells as in a and b with concentration as indicated. D) Orthotopic PSN1 (top) and PANC1 (below) tumors with representative H&E and Masson Trichrome staining with % stromal involvement in tumors, bars (including MiaPaca2, Supplementary Figure 8A). Mice harbouring established tumors were treated for 5 days with combined Metformin and AA and 5hmc and nuclear GATA6 quantified by IHC. E) RNAseq of orthotopic tumors with reads deconvoluted to examine Human tumor specific changes after treatment with Metformin and AA.

### Restoration of GATA6 in vivo is associated with a revival of WNT dependence

To determine if subtypes could be altered *in vivo*, mice harbouring PSN1, PANC1 and MiaPaca2 orthotopic tumors were treated with Metformin/AA. After five success days of treatment all tumors (distinguished by staining for human-specific mitochondrial membrane) displayed restoration of 5hmc and GATA6, implicating a tumor cell specific alteration towards a more classical-pancreatic identity, despite differential stromal involvement (Figure 6D and Supplementary Figure 7A). To verify if transcriptional patterns reflected a change in subtype, 3’RNA-seq of tumors were aligned to both the mouse and human genome and XenoFilter was used to deconvolute tumor vs stroma transcripts. GATA6 found to be increased in both tumor and stromal cells post treatment (S7B). This was again associated with a shift away from squamous identity and with a highly significant reduction in Wnt signatures (Wnt and β-catenin signalling) (Figure 6E and Supplementary Figure 8C). In PDAC tumors, stromal derived Wnt maintains the PDAC stem cell niche. However, tumors that lose GATA6 expression become Wnt independent by producing Wnt3A, Wnt7B and Wnt10A^33^. Elevated Wnt is linked to increased tumor aggressiveness of the squamous subtype as tumor cells are no longer reliant on the stromal support^44, 33^.

Metformin/AA resulted in a significant downregulation of WNT3A and R-Spondin 1 (RSPO1), and loss of nuclear β-catenin, implying that GATA6 correlates with Wnt production in classical-progenitor tumors (Figure 7A and Supplementary Figure 8A). Interestingly, cancer associated fibroblasts are also sensitive to 5hmc levels^45^ but decreased Wnt7b in stromal cells suggests that tumors cannot revert to a stromal dependent phenotype. Concomitantly, Metformin/AA treatment was sufficient to restrict tumor proliferation marker ki67, reduce capacity to invade *in vitro* and metastasise to the liver *in vivo* (Supplementary Figure 8B-D). Finally, to confirm the interaction with Wnt signalling we compared both GATA6 and TET2 expression with Wnt pathway genes, including RSPO1, Wnt3A, Wnt7B and Wnt10A in our patient cohort and found that expression of TET2 and the classical-pancreatic marker GATA6 are associated with reduced Wnt expression in human tumors (Figure 7B).

**Figure 7:**
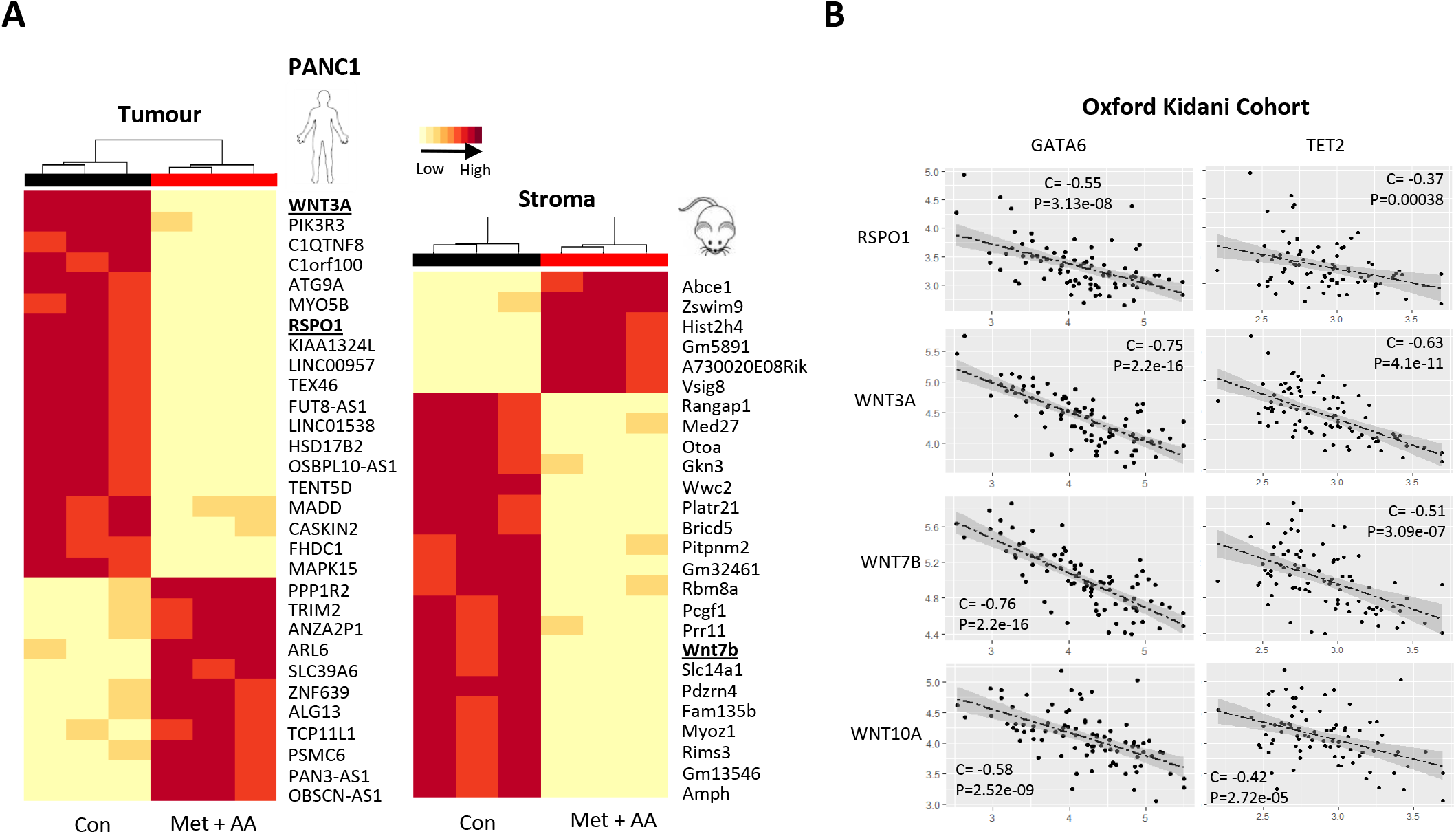
Restoration of 5hmc using Metformin and Ascorbic Acid Alters Subtype Identity. A) Differentially expressed genes in (tumor) and mouse (stroma). B) Correlation between *GATA6* and *TET2* mRNA with indicated Wnt ligands in Oxford-Kidani cohort.

## DISCUSSION

Transcriptomic subtypes define therapeutic outcomes in pancreatic cancer^5^. In retrospective clinical studies, classical-pancreatic tumors have been shown to be more therapeutically responsive and have 2-year survival of approximately 50%, whereas squamous-like subtypes have significantly poorer survival, reported at between 23 to 28% ‘. The therapeutic implication of these subtypes has now been demonstrated in a *prospective* feasibility trial (COMPASS,^5^), where patients with advanced PDAC (metastatic and locally advanced) underwent whole genomic characterisation identifying 70% patients with classical-pancreatic, and 30% with basal (squamous-like) subtype. Patients with classical-pancreatic subtype had statistically superior progression free survival (6.4 mo vs 2.3 mo, Hazard Ratio (HR) 0.28, p<0.001) and overall survival (10.4 v 6.3mo, HR 0.33, p<0.004) compared to patients with basal/squamous-like tumors^5^. GATA6 is a driver of PDX1 expression and pancreatic differentiation during development^46^and notably high tumor GATA6 was significantly correlated with the better differentiated classical-pancreatic subtype (p<0.001), thus proposed as a surrogate marker for differentiating subtypes^5^. Moreover, in human derived organoids, GATA6 alone was found responsible for transcriptional subtypes, as expression directly reverted squamous-like subtype to classical-progenitor, including the reversal of WNT-independent growth^33^, a signature we also find correlating with the epigenetic imbalance (Figure 3A). The fact that squamous-like tumors acquire a Wnt-autonomous / stromal-independent phenotype is proposed to permit greater dissemination and metastasis^33^, commensurate with more aggressive disease and poorer prognosis of this PDAC subtype. This raised the fundamental question of whether it may be possible to therapeutically switch pancreatic cancer subtype from squamous to classical by re-expressing GATA6, and therein potentially improve survival. Our novel epigenetic characterisation of the 5mc/5hmc methylome from FFPE sections of human tumors allowed us to identify 5hmc marking of GATA6 as a causative mechanism supporting expression of GATA6 itself and PDAC subtype. This suggested that manipulation of 5hmc may constitute a viable mechanism to reinstate GATA6 levels in squamous-like tumors. Moreover, the specific identification of a single 5hmc CpG correlating with GATA6 expression (cg21040686) may prove useful in the determination of transcriptional subtypes from circulating tumor-derived DNA in plasma^47^.

The TET-hydroxylases that are responsible for generation of 5hmc belong to a class of enzymes known as α-ketoglutarate-dependent enzymes and for which the α-ketoglutarate (α-KG) and Fe(II) are essential cofactors^8^. Recently, it was demonstrated that p53 promotes utilisation of glucose by the citric acid cycle, and as α-KG (also known as 2-oxoglutarate) is a central metabolite, maintaining sufficient levels of α-KG for TET enzymatic activity and 5hmc^48^. TET expression is supported by p53 during differentiation and concomitantly, when p53 is mutated the loss of 5hmc can contribute to PDAC progression in mouse models.^48, 49^. Here we find low levels of 5hmc in human tumors compared to normal pancreas, in agreement with high levels of p53 mutation, but an additional split in the tumor cohort based on epigenetic profiling that is associated with SMAD4 mutation. Importantly, SMAD4 mutation has been associated with more aggressive pancreatic cancers, and we demonstrate here that expression levels, rather than simply mutation, are closely associated with TET2 expression and levels of 5hmc. Mechanistically, we show that SMAD4 associates with the 3’ region of the TET2 gene in response to TGFβ signalling and confirm the positive correlation of SMAD4 with TET2 in our discovery cohort and the TCGA. Taken together, this implies that both p53 and SMAD4 are important for TET2 expression and, together with the co-factor α-KG, for 5hmc marking of genes. We propose that in the absence of p53, reduced activity of TET2 contributes to development of classical-pancreatic tumors, whereas the further loss of 5hmc through reduced TET2 expression results in conversion to squamous-like subtypes. This may involve a progression from classical-pancreatic to squamous-like subtype over time due to the influence of the tumor microenvironment or further metabolic stress, constituent with squamous being more apparent in later stage disease^50^.

While we find SMAD4 supports expression of TET2, this appears insufficient to sustain 5hmc in the absence of the TET2 co-factor α-KG^48^, emphasising how 5hmc levels are linked to metabolic regulation and the citric acid cycle. While α-KG-dependent hydroxylases, can be sensitive to oxygen levels, TET enzymes appear more refractory to low oxygen conditions and display increased activity under hypoxic conditions^51^. This suggests that the Fe(II)-dependent hydroxylation reaction that promotes 5mc-5hmc is supported by additional cofactors in hypoxia, e.g. such as AA. While not exactly clear how AA functions to support TET function, it is required to support α-KG and interestingly, developmental TET homologs have been described to directly modify nucleotides with AA^52^. Thus, supplementing AA or α-KG have both been independently demonstrated to support TET-mediated 5hmc, but obviously, this is dependent on levels of the enzymes itself. Notably, we observe that AA appears to stabilise TET2 protein level (Figure 6B), potentially through direct binding. However, the stability of TET2 is also susceptible to protein-turnover in a mechanism that responds to cellular energy via AMPK activation^40^. Interestingly, the anti-diabetic drug Metformin disrupts redox signalling to restore ATP levels and AMPK activity, which in turn stabilises TET2 as a direct substrate^40^. Therefore, as found for AA, Metformin also stabilises TET2 levels in PDAC, but alone each appears insufficient to restore 5hmc and expression of GATA6. However, combination of these treatments, at physiologically relevant doses to levels in human plasma, can restore TET2 levels, 5hmc, GATA6/PDX1 expression and transcriptional attributes of more differentiated tumors in vitro and in vivo, such as WNT-dependence. It is notable that in human tumors we find highly-significant inverse correlations between Rspondin1 coreceptor and WNT ligand expression (WNT3A,7B and 10) with both GATA6 and TET2 suggesting WNT autonomy in squamous-like tumors is linked to 5hmc and that supporting observation that classical-pancreatic tumors may be more dependent on stromal derived WNT signalling^33^. Critically, TET2 stability and 5hmc activity can be restored using a combination of Metformin and AA, resulting in GATA6 re-expression and a switch from squamous to classical-pancreatic identity in vivo.

Our work suggests it may be possible to therapeutically switch squamous-like PDAC to the more favourable classic-pancreatic subtype using Vitamin C and Metformin, thereby enhancing chemotherapy response and survival. Interestingly, as TET2 is sensitive to glucose levels via AMPK^40^, loss of glucose tolerance may also contribute to reduction of TET2 protein levels, loss of pancreatic identity and emergence of cancer in newly diagnosed pancreatic diabetes patients. Moreover, the factors contributing to loss of AMPK activity, e.g. reduced exercise or high fat diet^53^, together with reduced Vitamin C levels may help to explain the appearance of aggressive some pancreatic cancers. Taken together our results outline how the basis of transcriptional subtypes in cancer are defined through epigenetic regulation and how mutation and metabolic events promote a de-differentiation that is reversible *in vivo*.

## Methods

Supplementary Materials and Methods for full methods.

### Patients and Tissue

Formalin fixed paraffin embedded (FFPE) tissue blocks were obtained together with clinical follow-up data from the Oxford Radcliffe Biobank, Oxford University Hospital NHS Trust. Patients included in the present retrospective study had to meet the following criteria: histologically-confirmed PDAC, complete macroscopic surgical resection (R0 or R1), lack of metastatic spread and/or ascites, absence of previous history of malignancy and archived FFPE surgical samples at the Department of Pathology. In total, n = 141 patients were included in the total cohort.

### Oxidative Bisulphite sequencing (oxBS)

To determine genome-wide hydroxymethylation the CEGX TrueMethyl Array Kit was adapted for use with FFPE material with 1ug of DNA in 50ul ultrapure water used as starting material. DNA was first purified and denatured DNA was then split into two separate 0.2ml PCR tubes. The oxBS sample was oxidized by adding 1ul of Oxidant solution and incubating at 40 °C for 10 minutes. Following oxidation and centrifugation the sample should remain an orange colour (Supplementary Figure 2A), indicating excess oxidative reagent in the sample. All samples underwent bisulfite conversion by addition of 30ul bisulfite reagent and were then placed in a thermocycler. Tubes were centrifuged at 14,000g for 10 minutes and 40ul of supernatant was transferred to a new 1.5ml eppendorf. DNA was purified and resuspended in 200ul Desulfonation buffer, washed and DNA eluted in 12ul elution buffer. Samples were then run on illumina EPIC arrays by the Oxford Genomics Group, Oxford, UK.

### Genomics and sequencing

Isolation of mRNA from FFPE samples was performed using Qiagen RNeasy FFPE kits according to the standard protocol. Two 4um scrolls were used for each sample, and excess paraffin was removed by xylene washes. RNA was then eluted in 30ul RNase free water. RNA quality and quantity was determined by Agilent Bioanalyzer and Qubit Fluorometer (Thermo Fisher) respectively. Purified RNA was then run on Almac Xcel Arrays.

Genomic mutations in PDAC tumors were determined for a panel of 52 genes commonly mutated in multiple cancers. Sequencing libraries were generated using both the Oncomine BRCA Research Assay Chef-Ready kit (Life Technologies, Carlsbad, CA, USA), and the Ion Ampliseq Cancer Hotspot Panel V2 Chef-Ready kit (Life Technologies, Carlsbad, CA, USA). Only sequences with a p-value <0.001, sequencing depth >100x and a functional consequence (missense, nonsense or indel) were used. Average sequencing depth after filtering was 474.

### Statistics and bioinformatics

Unless stated otherwise, all statistical analysis was carried out using R version 3.4.3. In all figures: * = p < 0.05, ** = p < 0.005, *** = p < 0.0005 and ns = not significant. Students T-Tests were performed in Microsoft Excel for comparison of two groups. When multiple comparisons were made at once the Benjamini-Hochberg (BH) procedure was used with a false discovery rate of 10% to adjust for multiple comparisons. Chi-Square independence tests were used to compare categorical variables between groups, unless an expected value was less than 5, in which case Fisher’s exact tests were used. Correlation between two continuous variables was analysed using Pearson correlation coefficient. Kaplan-Meier curves and Cox Proportional Hazard Regressions were performed using the R Survival package. For IHC quantification of orthotopic tumors Mann-Whitney U Tests were performed in Graphpad Prism 7.0. Graphs were drawn using the ggplots2 package or Graphpad Prism 7.0, and clustered heatmaps drawn using the pheatmap package.

EPIC array data was processed using the R package ChAMP. Differential methylated probe (DMP) analysis was carried out using the champ.DMP() function using an adjusted P value cutoff of 0.05. Gene set enrichment of DMP/DhMPs was performed using the champ.GSEA() function. To determine the enrichment of probes at different genomic features the CpG.GUI function was used. To generate genome tracks for visualization of methylation/hydroxymethylation across genes, the average β-value at for each probe was taken for healthy and tumor samples or different molecular subtypes. These were then uploaded and visualized on the UCSC Genome Browser (http://genome/ucsc.edu/) assembly GRCH37/hg19. Gene sets were downloaded from the Broad Institute MSigDB database v6.2 (Hallmarks, c2 Curated, c5 Gene ontology and c6 oncogenic signatures) and enrichment of mRNA datasets or manually creating a gene set for each subtype performed using the R package GSVA. The Basal and Classical signatures from Moffit et al.^4^ for Collisson subtypes created using the genes listed in PDAssigner. Squamous, progenitor, ADEX and immunogenic gene sets were created using the core gene programmes listed in the ures by Bailey et al. (2016) (GP1, GP2, GP6 and GP9). GSVA results were hierarchically clustered using the pheatmaps package and subtypes were assigned according to which subtype geneset was enriched in each cluster. For determining molecular subtypes in cell lines, normalized ZScores from RNAseq of pancreatic cell lines was downloaded from the Cancer Cell Line Encyclopedia.

### iCluster

Integrated analysis of mutational, methylation and hydroxymethylation data was performed using the iClusterPlus package. For gene mutations, binary data was used (1 = mutated, 0 not-mutated) for all genes mutated in over 5% of samples. For methylation and hydroxymethylation data the top 5,000 most variable probes were used. Bayesian information criteria (BIC) modelling using the tune. iCluster function was used to determine the optimal number of clusters to explain the variation within the data. The probes and mutations most closely associated with each cluster were shown using the plotHeatmap function. Significant associations between clusters and mutations or clinical characteristics were determined by Chi-Squared or Fisher’s tests.

### Datasets

RNA-seq, Methylation 450K and Clinical data was downloaded for Pancreatic Adenocarcinoma (PAAD) by The Cancer Genome Atlas (TCGA) version 2016 1 28 from the Firebrowse portal (Firebrowse.org). A SMAD4 Chip-seq performed in A2780 cells was downloaded from the Gene Expression Omnibus (GEO) accession number GSE27526 and SMAD4 Tet-off and Tet-on mouse pancreatic cancer cell lines GSE72069.

### Orthotopic Tumors

Pancreatic orthotopic tumors were generated by injection of human pancreatic cell lines (PANC1, MiaPaca2 and PSN1) directly into the pancreata of athymic nude-Fox1nu mice. Tumor growth were monitored post-surgery using 4.7T MRI scan (Bruker BioSpin GmbH, Germany). In the treatment cohorts, Metformin (Abcam) was provided in the drinking water (2.5mg/ml) from day 1 (11 days after the surgery). Sodium Ascorbate (Sigma-Aldrich) was administrated (2g/kg in PBS) via intraperitoneal injection daily from day 1 until day 5 concurrently with Metformin. Mice were euthanized at day 5 of the treatment.

DNA, RNA and miRNAs were simultaneously purified from orthotopic tumors using Qiagen AllPrep DNA/RNA/miRNA Universal Kits. Tissue was disrupted using TaKaRa biomashers and homogenized using Qiagen QIAshredders. Quantity and quality of DNA and RNA was determined using Nanodrop, and RNA integrity was analyzed using Agilent Bioanalyzer. Paired 3’RNA-seq was performed on isolated RNA by the Oxford Genomic Centre, Oxford, UK. Following sequencing, raw fastq reads were aligned to the human (hg19) and mouse (mm9) genomes using RNA STAR on the Galaxy web platform. To deconvolute incorrectly aligned mouse/human reads the R package XenoFilter was used (Kluin et al., 2018). Following deconvolution, a count matrix was generated using the GenomicFeatures and GenomicAlignments packages and processed using the DESeq2 package.21

## Authors contributions

SM and EO’N conceived the project. ME, SL and E’ON designed experiments. ME, SL and FW performed all experiments, ABu and AS performed DNA array, AB, DH and DV assisted in database building and data analysis, RG performed pathology, ME and ZDC optimised the FFPE-oxBS protocol. AH, ZDC, AGA, AS, SJ and CV performed sample collection and followup data. ZS and MS performed all resections. ME, AB and TM assisted with RNA analysis ME and EON wrote the manuscript. SM and EON supervised the study.

## Acknowledgements

This work was funded by the Kidani Memorial Trust, Cancer Research UK Oxford Cancer centre, Cancer Research UK Precison Panc consortium and a Medical Research Council studentship (ME). Authors are grateful to H.Kosher for hTert-DEC cells and E.Fokas, and M.Youdell for invaluable assistance in sample collection.

---

**Supplementary Figure 1:**
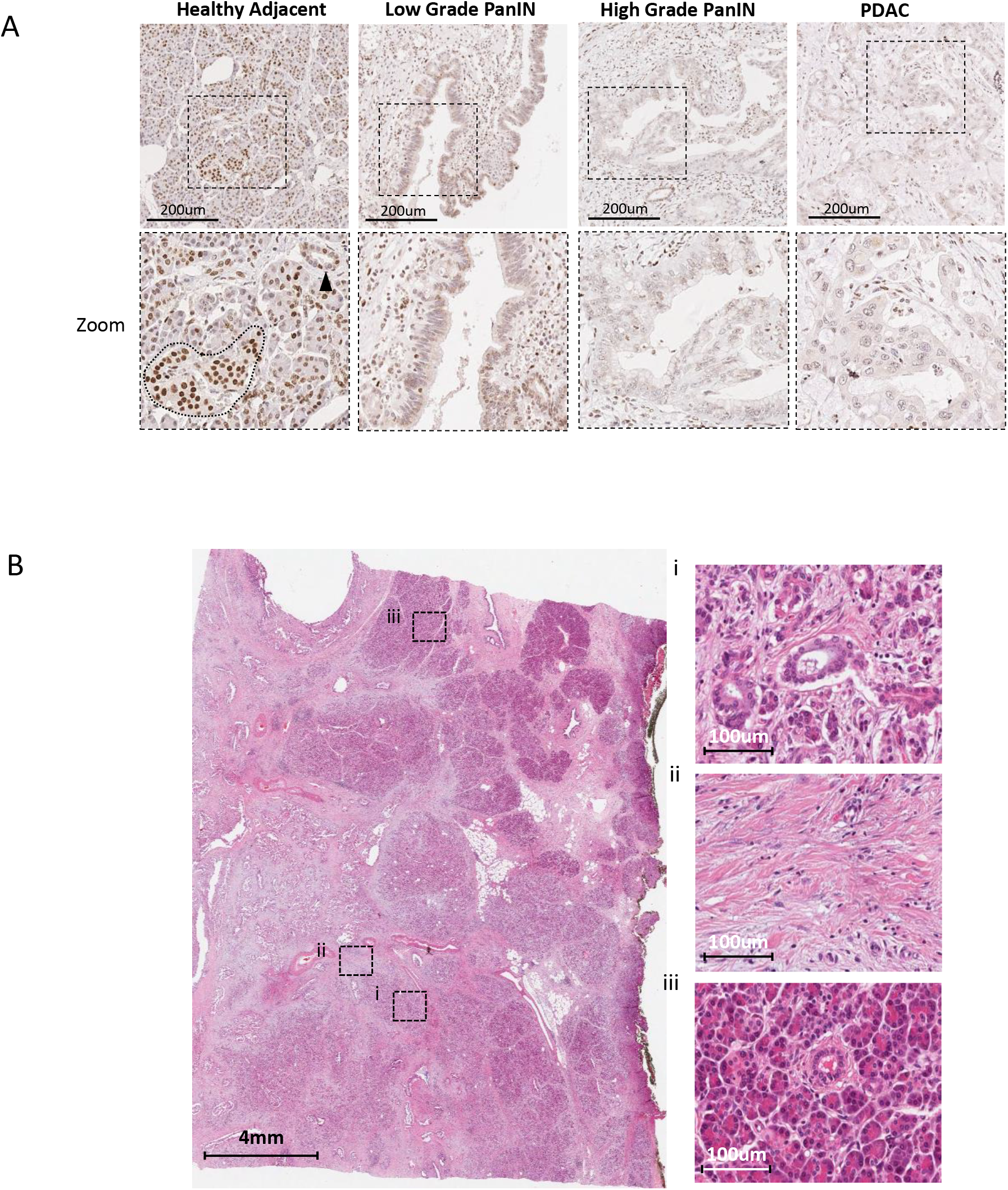
A) IHC for 5hmC in human pancreatectomy sections. 5hmC staining intensity was quantified in healthy adjacent tissue, low and high grade PanINs and neoplastic cells (PDAC). 5hmC staining is progressively lost during PDAC progression. B) Example H&E of whole pancreatectomy sections used for analysis. Right, regions of (i) tumour, (ii) stromal and (iii) healthy adjacent tissue within the sections.

**Supplementary Figure 2:**
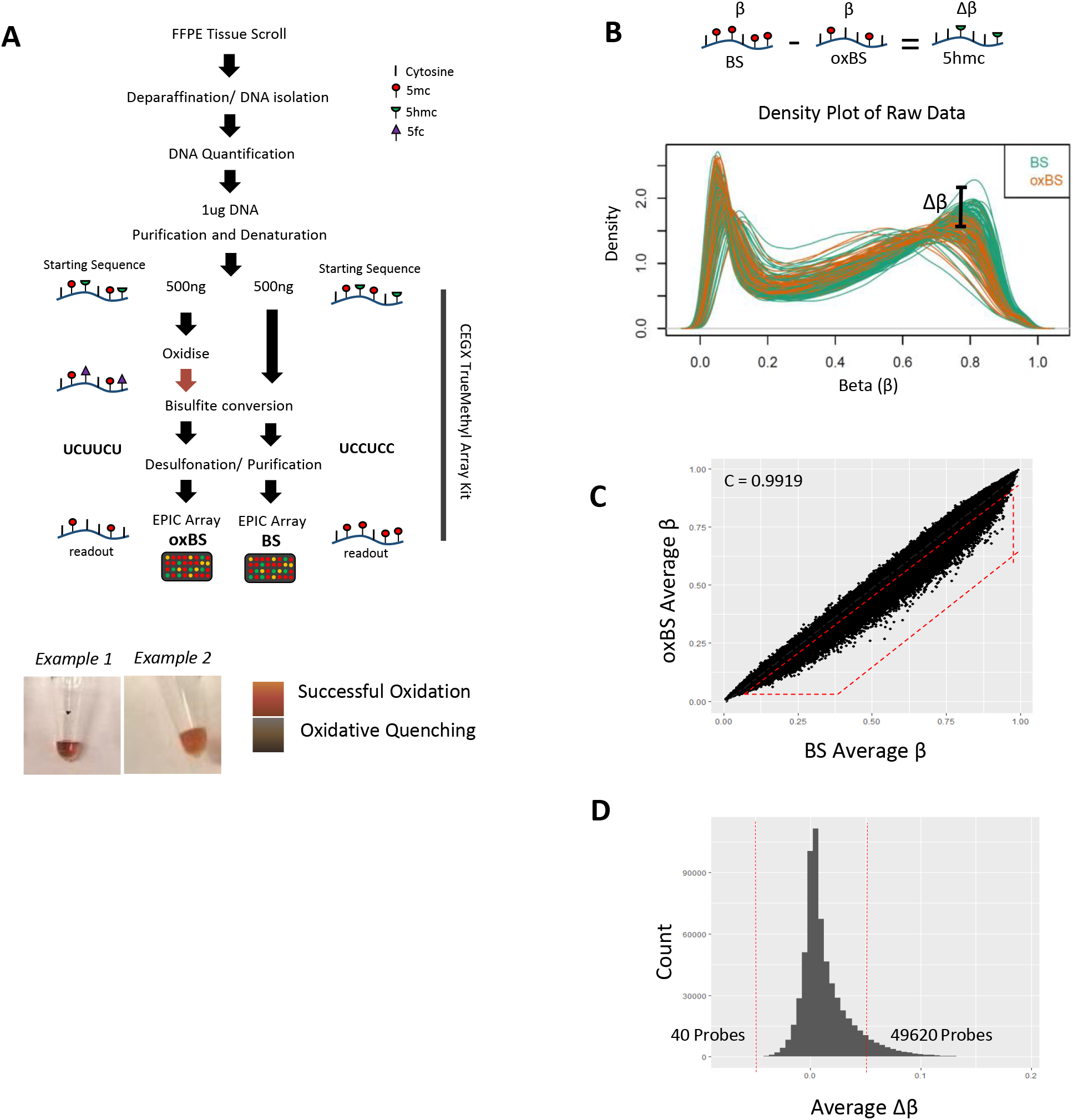
Adaption of the oxBS method to FFPE. **A)** Work flow for oxBS method. DNA is split into two 500ng samples, one of which undergoes conventional bisulfite treatment (BS), while the other is oxidized prior to bisulfite treatment (oxBS). below: Examples of successful oxidation. The orange colour indicates residual Potassium Perruthenate oxidative reagent, meaning the reaction has not been quenched. 5hmC levels are determine by subtracting oxBS from BS values. **B)** Raw density plots from EPIC arrays for BS and oxBS samples. Oxidation of 5hmC results in a drop-in signal for oxBS samples. **C)** Correlation between average probe β-values in BS and oxBS samples. Lower signal in oxBS sample relative to BS is due to loss of 5hmC signal (red box). **D)** Average Δβ across all probes. Positive Δβ values indicate hydroxymethylated probes, while negative Δβ values are due to background noise. Only 40 probes have Δβ below −0.05, while almost 50,000 probes had Δβ values above 0.05. E) Scatterplot showing correlation between 5hmC at CpG shores and 5mC within CpG islands in healthy and tumour samples shows a highly significant negative correlation.

**Supplementary Figure 3:**
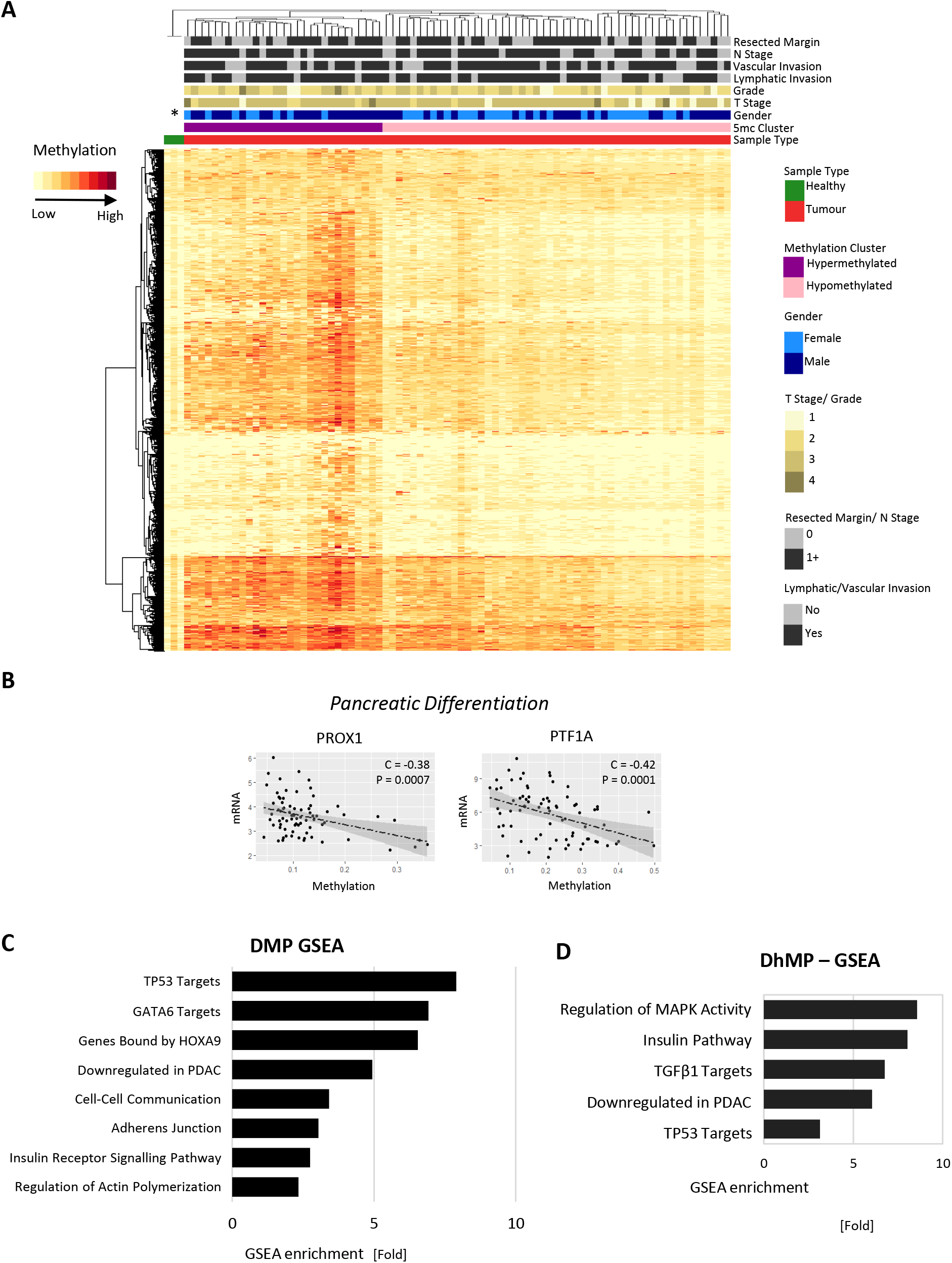
Methylome of Oxford Kidani Cohort Tumours. **A)** Differentially methylated probes clustering. **B)** GSEA of enriched DMPs. **C)** Methylation negatively correlates with mRNA at pancreatic differentiation genes. All methylation analysis was performed using oxBS data.

**Supplementary Figure 4:**
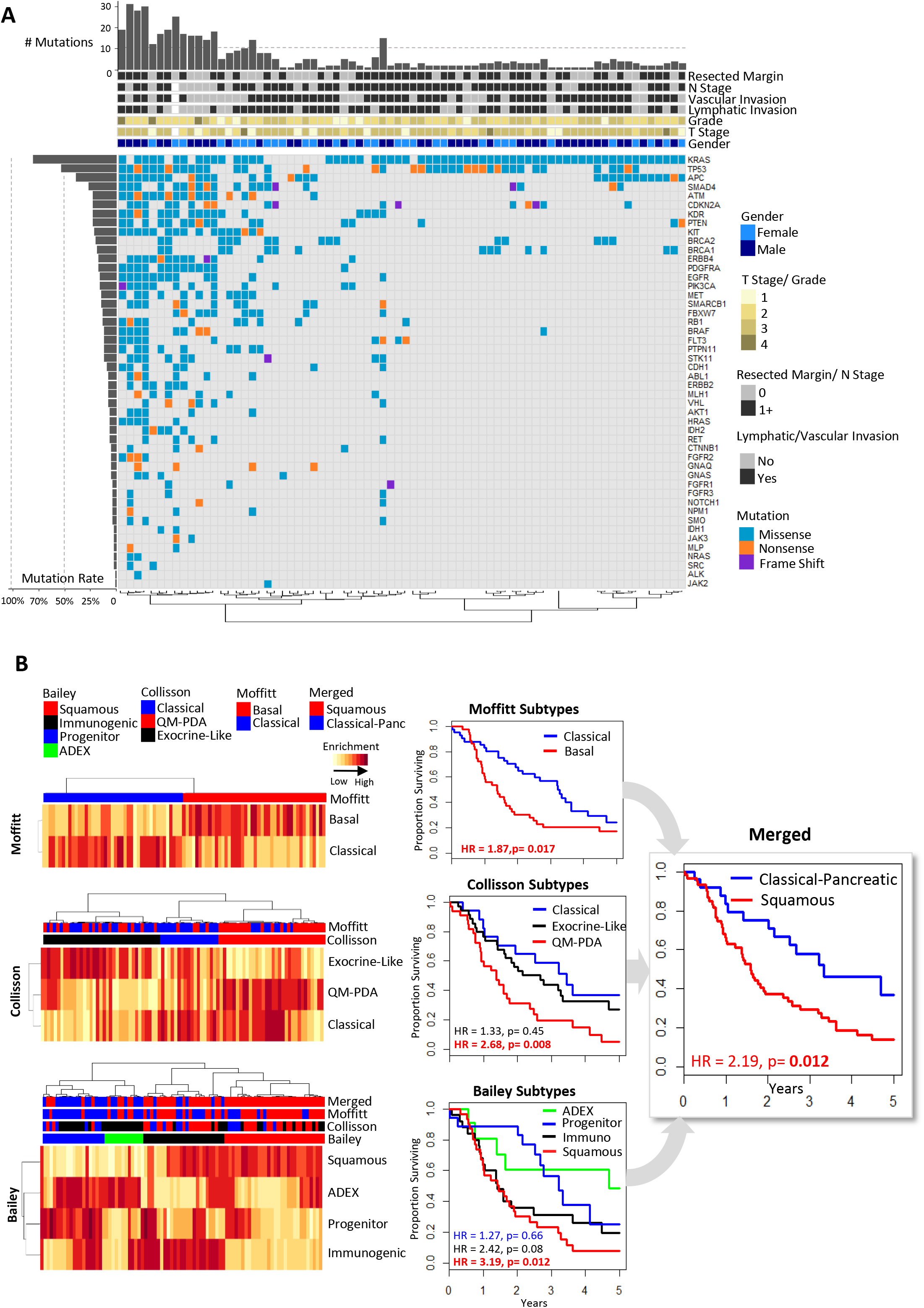
Molecular Subtyping of the Oxford Kidani Cohort. A) Overview of focussed mutational profiling of pancreatic tumours. Top panel shows the individual mutation rates per patient from the 52-gene panel. Middle panel shows clinical characteristics. Left panel shows the mutation rates per gene. Genes in which no mutations were detected are not shown. KRAS, TP53 (including the germline p.P72R associated with pancreatic dysplasia, Kung et al. Cell Reports 2016), SMAD4 and CDKN2A mutation rates are in line with previous PDAC sequencing studies. APC mutations include somatic missense mutations not documented to be pathogenic. B) Molecular subtyping of tumours using defined gene sets1-3. Relative enrichment of each gene set was determined using GSVA. Subtype for each patient is shown in the top panel of each heatmap. Right) Kaplan-Meier curves for patients of each subtype. In all cases patients with the more aggressive subtype (Basal, QM-PDA and Squamous) had a worse prognosis. C) Kaplan-Meier survival analysis for merged squamous and classical-pancreatic subtypes.

**Supplementary Figure 5:**
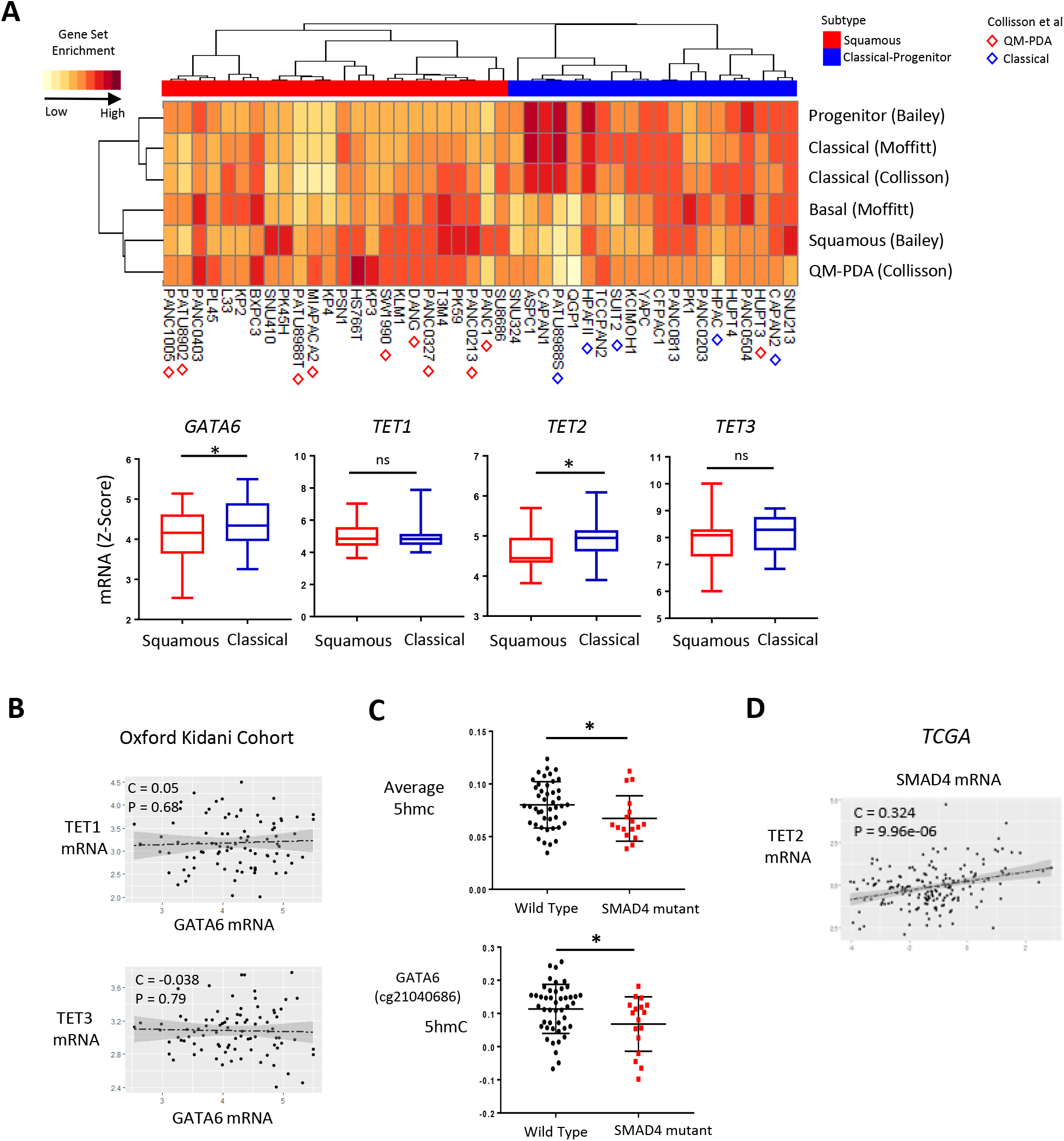
Molecular Subtyping of PDAC cell lines. A) Subtyping of CCLE PDAC cell lines based on previous studeis^1–3^. Enrichment for subtype gene sets was determined using GSVA. Results were hierarchically clustered and cells were assigned a subtype of Squamous or Classical-pancreatic. ADEX, Exocrine-like and immunogenic subtypes were not included as they have not been found to be represented in cell lines. Red/blue triangles indicate cell lines classified by Collisson et al.3 Below, box and whisker plot showing differences in GATA6, TET1, TET2 and TET3 mRNA for squamous and classical-pancreatic cell lines from CCLE data. Squamous cell lines have lower levels of GATA6 and TET2. B) Correlations between TET1 and TET3 with GATA6 mRNA in the Oxford Kidani cohort. C) (top) Total DhMPs and (bottom) GATA6 cg21040686 in tumours with wild type and mutant SMAD4. D) Correlation between TET2 and SMAD4 in the TCGA_PAAD cohort.

**Supplementary Figure 6:**
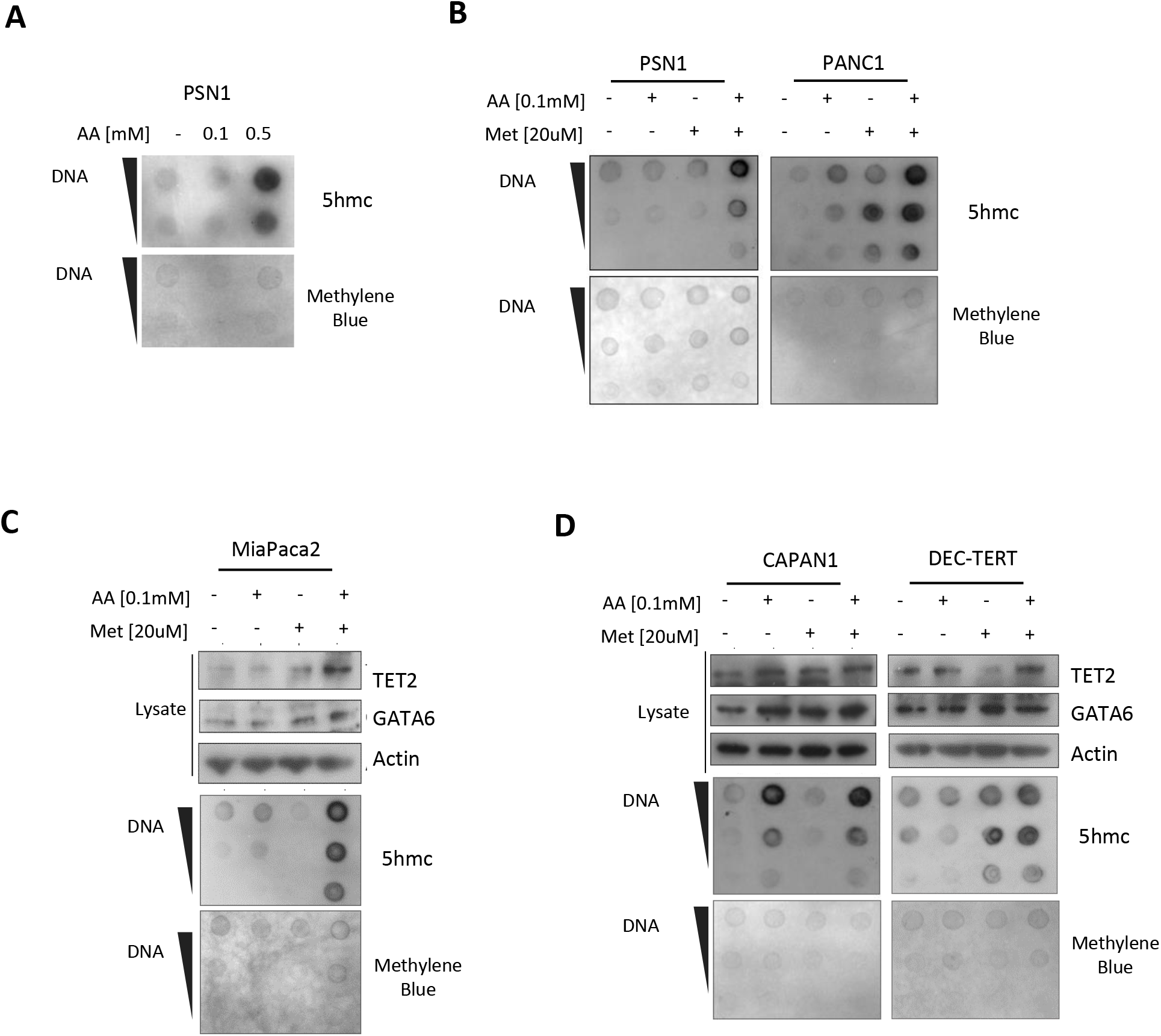
Metformin and Ascorbic Acid Restore 5hmC and GATA6 in Squamous cell lines. A) Dot blot dilutions for PSN1 cells were treated with AA or (B) combined Metformin and AA represented in Fig. 3b,c. Western blots and dot blots of (C) MiaPaca2, (D) CAPAN1 and DEC-hTERT cell lysates after treatment with indicated concentrations of Metformin and AA.

**Supplementary Figure 7:**
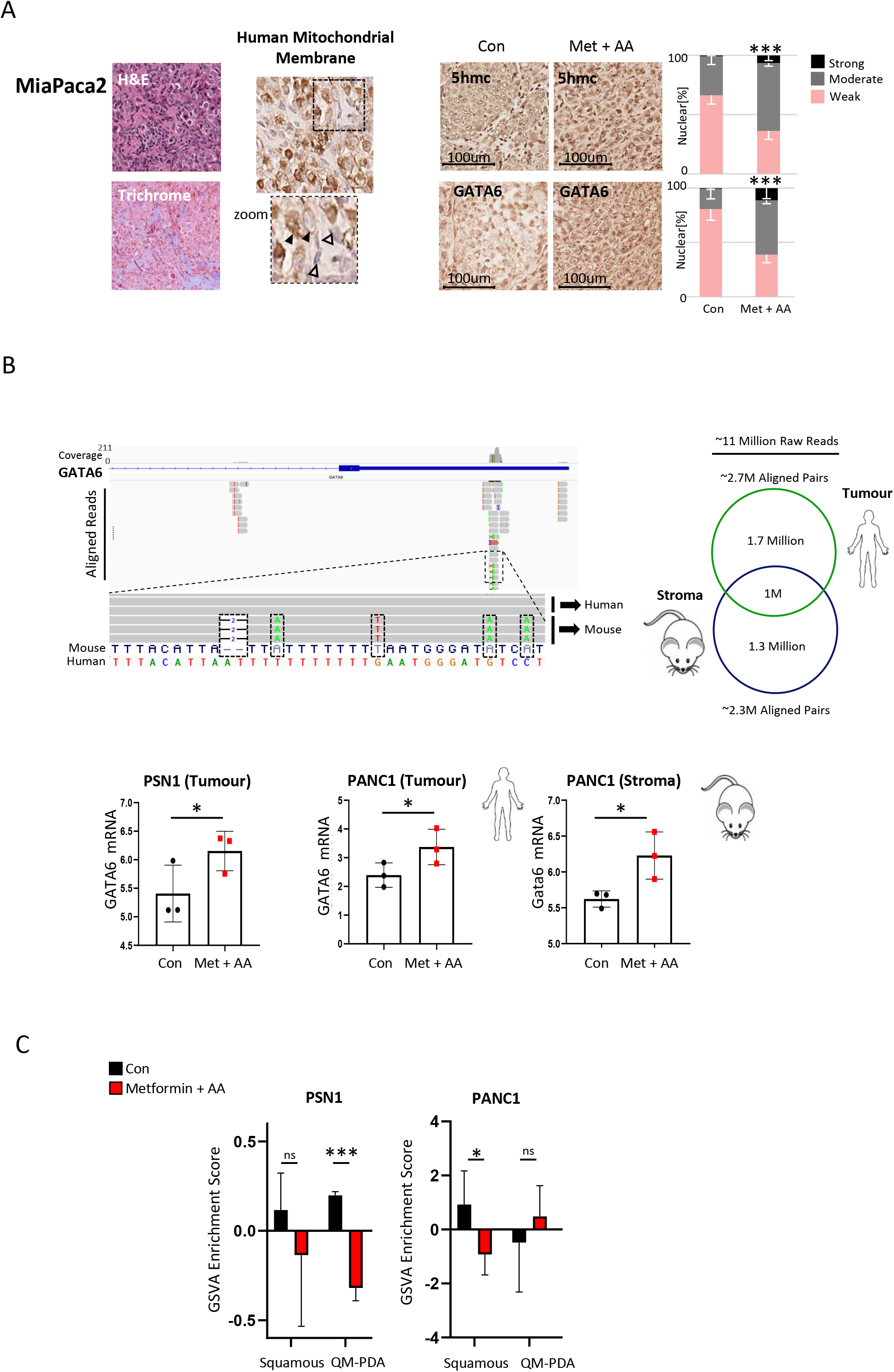
RNA-seq of orthotopic tumours indicates partial change in molecular subtype. A) Orthotopic MiaPaca2 tumours with representative H&E and Masson Trichrome staining. Mice harbouring established tumours were treated for 5 days with combined metformin and AA and 5hmC and nuclear GATA6 quantified by IHC. B) top, IGV genome viewer showing RNA-seq reads aligned to the human genome (hg19). Missmatch reads perfectly correspond to the mouse sequence for the same region, indicating incorrectly aligned mouse reads. Mouse and human reads were deconvoluted using the XenoFilter programme. Below, changes in GATA6 mRNA following Metformin/AA in PSN1, PANC1 tumours and PANC1 tumour associated mouse stromal cells. RNA-seq for PSN1 stroma cells could not be determined due to the lower proportion of mouse cells in these tumours. C) Enrichment of molecular subtype gene sets following treatment of PSN1 and PANC1 orthotopic tumours.

**Supplementary Figure 8:**
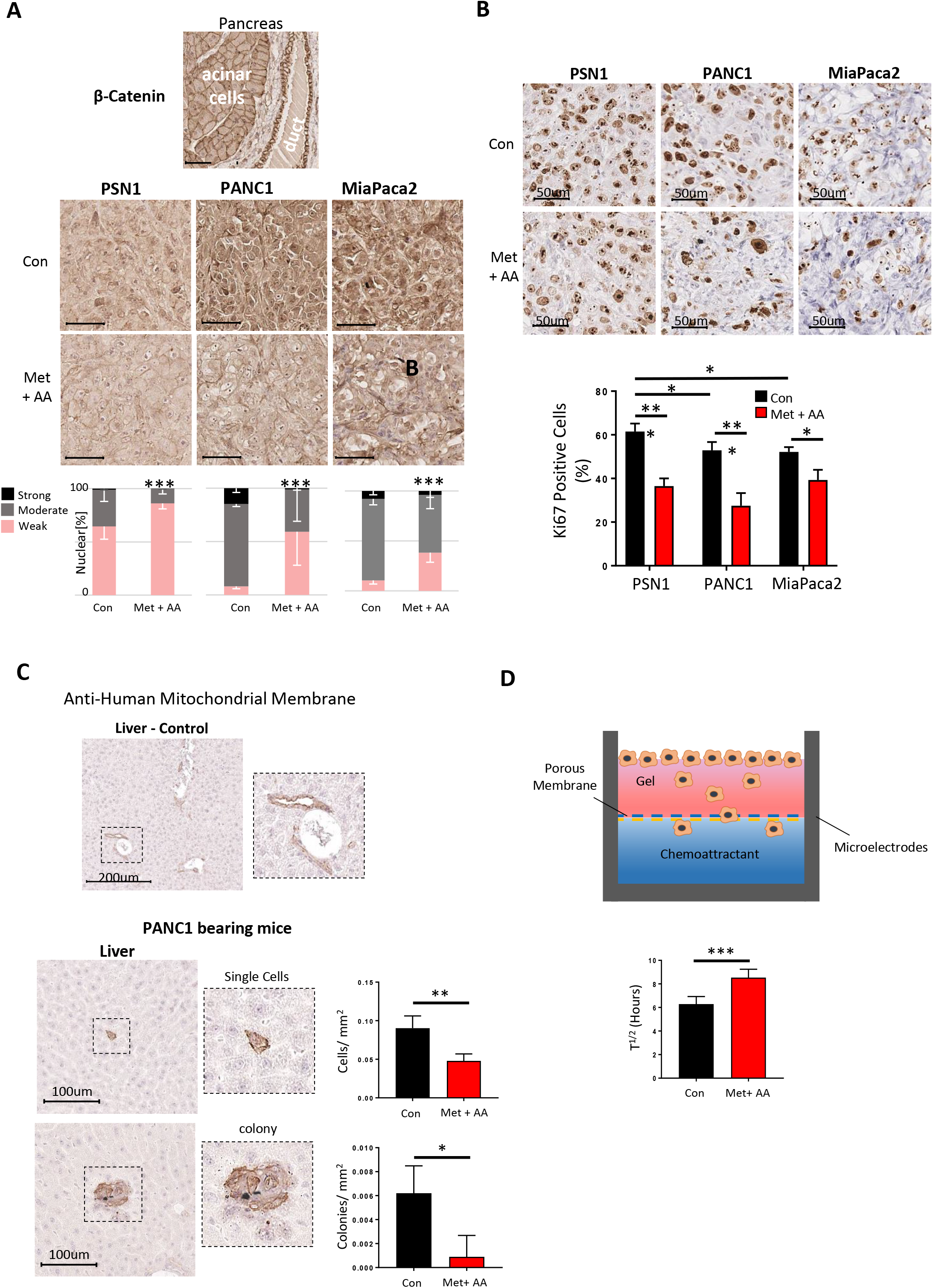
Metformin/AA treatment inhibits nuclear β-catenin. **A)** Top: IHC of healthy mouse pancreatic tissue shows membranous β-catenin in acinar and ductal cells. below) β-catenin is nuclear in orthotopic tumours. Following treatment nuclear β-catenin is reduced. **B)** ki67 staining of orthotopic tumours shows inhibition of cell proliferation following treatment. **C)** Identification of disseminated tumour cells in livers of tumour bearing mice with a specific antibody for human mitochondrial membranes and distinguished by nuclei shape (large irregular human tumour cells vs small oval mouse stroma cells). For IHC analysis only human cells were quantified. Top: negative control showing lack of staining throughout the liver on non-tumour bearing mice. Some weak background staining can be seen lining duct cells. Below: IHC staining of livers of PANC1 tumour bearing mice and quantification of micrometastatic individual cells and small clusters of tumour cells of Metformin/AA treated and untreated mice. D) Transwell migration assay of PANC1 cells. Metformin/AA treatment inhibited migration of cells *in vitro*.

**Supplementary Table 1:** Clinicopathological features of the cohort and association with overall survival. Significant p values in bold.

| Clinicopathological Feature | % | Hazard Ratio | 95% CI | P Value |
| --- | --- | --- | --- | --- |
| **Gender** |  |  |  |  |
| Female | 0.46 | 1 |  |  |
| Male | 0.54 | 0.835 | 0.573-1.217 | 0.349 |
| **T Stage** |  |  |  |  |
| T1 | 0.09 | 1 |  |  |
| T2 | 0.13 | 2.421 | 0.919-6.375 | 0.073 |
| T3 | 0.72 | 3.252 | 1.418-7.458 | **0.005** |
| T4 | 0.06 | 3.412 | 1.096-10.63 | **0.034** |
| **Grade** |  |  |  |  |
| G1 | 0.06 | 1 |  |  |
| G2 | 0.56 | 1.04559 | 0.475-2.29 | 0.912 |
| G3 | 0.36 | 1.83234 | 0.823-4.07 | 0.138 |
| G4 | 0.02 | 0.60976 | 0.126-2.93 | 0.537 |
| **N Stage** |  |  |  |  |
| N0 | 0.23 | 1 |  |  |
| N1+ | 0.77 | 2.632 | 1.563-4.433 | **0.0002** |
| **Resection Margin** |  |  |  |  |
| R0 | 0.38 | 1 |  |  |
| R1+ | 0.62 | 1.911 | 1.271-2.874 | **0.002** |
| **Lymphatic Invasion** |  |  |  |  |
| No | 0.24 | 1 |  |  |
| Yes | 0.76 | 2.2593 | 1.372-3.721 | **0.0014** |
| **Venous Invasion** |  |  |  |  |
| No | 0.32 | 1 |  |  |
| Yes | 0.68 | 2.0973 | 1.367-3.217 | **0.0007** |
| **Recurrence** |  |  |  |  |
| No | 0.34 | 1 |  |  |
| Yes | 0.66 | 2.8376 | 1.774-4.54 | **0.00001** |
| **Site of Recurrence** |  |  |  |  |
| Local | 0.21 | 1 |  |  |
| Distant | 0.29 | 1.5545 | 0.8242-2.932 | 0.173 |
| Both | 0.50 | 2.5616 | 1.4292-4.591 | **0.0016** |

## Supplementary Methods

### Patients and Tissue

Formalin fixed paraffin embedded (FFPE) tissue blocks were obtained together with clinical follow-up data from the Oxford Radcliffe Biobank, University of Oxford (REC approval from South Central C, 09/H0606/5+5; OCHRe Project ref: 14/A176) at the Oxford University Hospital NHS Trust. Patients included in the present retrospective study had to meet the following criteria: histologically-confirmed PDAC, complete macroscopic surgical resection (R0 or R1), lack of metastatic spread and/or ascites, absence of previous history of malignancy and archived FFPE surgical samples at the Department of Pathology. In total, n = 141 patients were included in the total cohort. Most patients were treated with adjuvant chemotherapy consisted of gemcitabine (GEM) as monotherapy at the Oxford University Hospital NHS Trust, Oxford, UK. GEM alone was administered intravenously (dose 1,000 mg/m2 over 30 minutes), at days 1, 8 and 15 (1 cycle) for up to 6 cycles. A few patients received a combination of gemcitabine with capecitabine (GEM-CAP). In this case, GEM was administered as described above and CAP was administered orally (dose 830 mg/m2 twice daily) for 3 weeks followed by 1-week pause.

### Histological Analysis

4um FFPE Sections on glass slides were de-waxed in two washes of Citroclear and rehydrated in graded alcohol concentrations before washing in PBS. For Haematoxylin and Eosin (H&E) staining slides were placed in freshly filtered Harris Haematoxylin for 2 minutes, differentiated in 1% acid alcohol solution and counterstained in eosin solution for 1 minute. For immunohistochemistry (IHC) antigen retrieval was performed in Tris-EDTA buffer (pH 9.0) at 125C in a pressure cooker. Sections were blocked using Dako envision+ Dual Endogenous Enzyme Block for 1 hour before incubating with primary antibody overnight at 4C. When a mouse antibody was used on mouse tissue an additional blocking step was performed using Mouse on Mouse Blocking Reagent (Vector Labs). Antibodies and dilutions used were 5’hmC (Active Motif, 1:100), GATA6 (Abcam, 1:100). Human Mitochondrial Membrane (Merck, 1:100), ki67 (Merck, 1:1000) and β-catenin (Santa Cruz, 1:100). Slides were washed with PBS before incubating with secondary antibody-HRP (Dako) for 30 minutes at room temperatures and then washed in PBS again. Slides were developed with 3,3’-diaminobenzidine (DAB) solution (Dako) and inactivated in water before counter staining in haematoxylin for 30 seconds. After all staining protocols slides were dehydrated in graded alcohol concentrations before being cleared in xylene washes and mounted in Di-n-Butyl phthalate in Xylene (DPX) and scanned using an Aperio ScanScope slide scanner. Human PDAC tissue was quantified by two independent pathologists. Orthotopic tumours were quantified using ImageJ. Only staining in the nuclei of tumour cells was quantified using the criteria that cells with small oval nuclei were stromal cells of mouse origin.

### Oxidative Bisulphite sequencing (oxBS)

In order to determine genome wide hydroxymethylation the CEGX TrueMethyl Array Kit was adapted for use with FFPE material. 1ug of DNA in 50ul ultrapure water was used as starting material. The use of high-quality DNA without ethanol or paraffin contaminants was found to be critical for successful completion of the protocol. Low retention pipette tips were used throughout to prevent loss of magnetic beads. DNA was extracted using Qiagen QIAamp DNA FFPE tissue kits with Qiagen Deparaffination solution. 1ug of DNA in 50ul ultrapure water was used as starting material. The use of high quality DNA without ethanol or paraffin contaminants was found to be critical for successful completion of the protocol. Additionally, all steps prior to sample oxidation were performed in a room where ethanol had not been sprayed for at least 12 hours to further reduce the risk of ethanol contamination. Low retention pipette tips were used throughout to prevent loss of magnetic beads. DNA was first purified by adding 100ul of Magnetic Bead Binding Solution 1 (MBBS1) and incubating for 20 minutes at room temperature. Tubes were then placed on a magnetic rack for 5 minutes to immobilize magnetic beads and the supernatant removed. Beads were washed and aspirated 3 times in 200ul 80% acetonitrile in a fume hood. After the final wash excess acetonitrile was aspirated and pellets were left to air dry for 5 minutes. DNA was denatured by adding 20ul of denaturing solution directly to the bead pellet. Samples were vortexed to ensure proper resuspension of beads and then incubated for 5 minutes at 37 °C before separating on a magnetic rack again. The supernatant containing denatured DNA was then split into two separate 0.2ml PCR tubes. One tube was labelled as the oxBS sample, and the other the BS sample. The oxidative step was carried out on the oxBS sample only. The oxBS sample was oxidized by adding 1ul of Oxidant solution and incubating at 40 °C for 10 minutes. The sample was then centrifuged at 14,000g for 10 minutes to pellet any precipitate that formed during oxidation and the supernatant transferred to a new tube. Following oxidation and centrifugation the sample should remain an orange colour (Supplementary Figure 2A), indicating excess oxidative reagent in the sample.

All samples underwent bisulfite conversion by addition of 30ul bisulfite reagent and were then placed in a thermocycler. DNA was denatured at 95°C for 5 minutes, incubated at 60°C for 20 minutes, denatured at 95°C for 5 minutes, incubated at 60°C for 40 minutes, denatured at 95°C for 5 minutes and then incubated at 60°C for 45 minutes before holding at 20°C. Tubes were centrifuged at 14,000g for 10 minutes and 40ul of supernatant was transferred to a new 1.5ml eppendorf. DNA was purified by adding 160ul Magnetic Bead Binding Solution 2 (MBBS2) and incubating for 5 minutes. Samples were then placed on magnetic racks for 15 minutes to allow beads to separate. The supernatant was discarded and the pellets washed in 200ul of 70% ethanol. The bead pellet was resuspended in 200ul Desulfonation buffer for 5 mins before placing on the magnetic rack again, aspirating the supernatant, and washing with 200ul 70% ethanol 3 times. After washing and aspirating the bead pellet was allowed to air dry for 15 minutes. DNA was the eluted in 12ul elution buffer. Beads were incubated for 10 mins at room temperature before being separated on a magnetic rack and the supernatant transferred to a new 1.5ml tube. Samples were then run on illumina EPIC arrays by the Oxford Genomics Group, Oxford, UK.

### Ion Torrent Sequencing

Genomic mutations in PDAC tumours were determined for a panel of 52 genes commonly mutated in multiple cancers. Sequencing libraries were generated using both the Oncomine BRCA Research Assay Chef-Ready kit (Life Technologies, Carlsbad, CA, USA), and the Ion Ampliseq Cancer Hotspot Panel V2 Chef-Ready kit (Life Technologies, Carlsbad, CA, USA). Libraries were barcoded on an Ion Chef instrument (Life Technologies) and sequenced on an Ion Torrent PGM System using Ion 318 sequencing chips (Life Technologies). Data analysis was performed using Torrent Suite Software V.5.2. Only sequences with a p-value <0.001, sequencing depth >100x and a functional consequence (missense, nonsense or indel) were used. Average sequencing depth after filtering was 474. Mutation rates in KRAS (78%), TP53 (50%), SMAD4 (26%) and CDKN2A (22%) were at similar rates to previous sequencing studies on PDAC tumours.

### Statistics and bioinformatics

Unless stated otherwise, all statistical analysis was carried out using R version 3.4.3. In all figures: * = p < 0.05, ** = p < 0.005, *** = p < 0.0005 and ns = not significant. Students T-Tests were performed in Microsoft Excel for comparison of two groups. When multiple comparisons were made at once the Benjamini-Hochberg (BH) procedure was used with a false discovery rate of 10% to adjust for multiple comparisons. Chi-Square independence tests were used to compare categorical variables between groups, unless an expected value was less than 5, in which case Fisher’s exact tests were used. Correlation between two continuous variables was analysed using Pearson correlation coefficient. Kaplan-Meier curves and Cox Proportional Hazard Regressions were performed using the R Survival package. For IHC quantification of orthotopic tumours Mann-Whitney U Tests were performed in Graphpad Prism 7.0. Graphs were drawn using the ggplots2 package or Graphpad Prism 7.0, and clustered heatmaps drawn using the pheatmap package.

EPIC array data was processed using the R package ChAMP(1). Probes with a low detection p-value were removed and samples with 5% of probes with low detection were excluded from analysis. Probes on the XY chromosomes, nonCG probes, those identified as containing SNPs and probes that align to multiple locations were filtered, leaving 625,994 probes for analysis. Differential methylated probe (DMP) analysis was carried out using the champ.DMP() function using an adjusted P value cutoff of 0.05. Due to the presence of negative β-values in the hydroxymethylation dataset, differential hydroxymethylated probe (DhMP) analysis was performed manually by two-tailed t-test followed by BH adjustment for multiple comparisons. Gene set enrichment of DMP/DhMPs was performed using the champ.GSEA() function. To determine the enrichment of probes at different genomic features the CpG.GUI function was used. The distribution of DMP/DhMPs was then compared to the distribution of all probes using chi-squared test. To generate genome tracks for visualization of methylation/hydroxymethylation across genes, the average β-value at for each probe was taken for healthy and tumour samples or different molecular subtypes. These were then uploaded and visualized on the UCSC Genome Browser (http://genome/ucsc.edu/) assembly GRCH37/hg19.

Gene set enrichment of mRNA datasets was performed using the R package GSVA, which can determine the relative enrichment of a gene set across a dataset(2). Gene sets were downloaded from the Broad Institute MSigDB database v6.2 (Hallmarks, c2 Curated, c5 Gene ontology and c6 oncogenic signatures). For determining molecular subtypes GSVA was also used by manually creating a gene set for each subtype. The Basal and Classical signatures from Moffitt et al.(2,3) were taken from Figure 3A and Gene sets for Collisson subtypes were created using the genes listed in PDAssigner in Figure 1. Squamous, progenitor, ADEX and immunogenic gene sets were created using the core gene programmes listed in the supplementary figures by Bailey et al. (4) (GP1, GP2, GP6 and GP9). The relative enrichment of each subtypes gene set was then calculated. For each set of subtypes the GSVA results were hierarchically clustered using the pheatmaps package and subtypes were assigned according to which subtype geneset was enriched in each cluster. For determining molecular subtypes in cell lines, normalized ZScores from RNAseq of pancreatic cell lines was downloaded from the Cancer Cell Line Encyclopedia(5). As Exocrine-like and ADEX subtypes have not been reported in cell lines and immunogenic subtypes are dependent on immune cells within the tumour microenvironment these subtypes were excluded from the cell line analysis. The remaining subtype enrichment scores were hierarchically clustered and cell lines assigned a squamous or classical-pancreatic subtype depending on which gene sets were enriched in each cluster.

### iCluster

Integrated analysis of mutational, methylation and hydroxymethylation data was performed using the iClusterPlus package(6). For gene mutations, binary data was used (1 = mutated, 0 not-mutated) for all genes mutated in over 5% of samples. For methylation and hydroxymethylation data the top 5,000 most variable probes were used. Bayesian information criteria (BIC) modelling using the tune.iCluster function was used to determine the optimal number of clusters to explain the variation within the data. The probes and mutations most closely associated with each cluster were shown using the plotHeatmap function. Significant associations between clusters and mutations or clinical characteristics were determined by Chi-Squared or Fisher’s tests.

### Datasets

RNA-seq, Methylation 450K and Clinical data was downloaded for Pancreatic Adenocarcinoma (PAAD) by The Cancer Genome Atlas (TCGA) version 2016 1 28 from the Firebrowse portal (Firebrowse.org). A SMAD4 Chip-seq performed in A2780 cells was downloaded from the Gene Expression Omnibus (GEO) accession number GSE27526. Expression data for SMAD4 Tet-off and Tet-on mouse pancreatic cancer cell lines by David et al. (7) was downloaded from GSE72069.

### Cell lines

PANC1, PSN1, MiaPaca2, CAPAN1 (originally obtained from ATCC and authenticated by short tandem repeat profiling) and human Ductal Epithelial Cells immortalised with telomerase, DEC-hTERT^1^ were maintained in Dulbecco’s Modified Eagle Medium (DMEM) supplemented with 10% (v/v) Fetal Bovine Serum (FBS), 2 mM Glutamine and 100 U/mL Penicillin/Streptomycin (Pen/Strep). Cells were maintained at 37C, with 5% CO2 in a humidified incubator. For in vitro drug treatments of cell lines, PANC1, MiaPaca2 and Capan1 cells were treated for 96 hours, and PSN1 and DEC-hTERT cells were treated for 48 hours due to faster doubling time. Complete DMEM supplemented with L-Ascorbic Acid (Sigma, A4403) and/or Metformin (Abcam, ab146725) were prepared fresh in dH2O or PBS respectively for each day of treatment. Compound C (Santa Cruz, sc-200689) was dissolved in DMSO at a stock solution of 20mM and stored at −20 °C. All drugs used were dissolved in the appropriate buffer and filtered through a 0.4um filter syringe unit to sterilize prior to use. Following treatment cells were harvested and protein, DNA and RNA were isolated for western blot, dot blot and qPCR analysis.

All transfection of cell lines was performed using Lipofectamine 2000 (Life Technologies) following the manufacturers guidelines. A ratio of 1ug siRNA or 1ug plasmid DNA to 1uL Lipofectamine was used. Cells were harvested 48 hours after transfection. For siRNA experiments siLuciferase was used as a control (Eurofins, A4403, GCCAUUCUAUCCUCUAGAGGAUG). Human SMAD4 siRNA was purchased from Dharmacon (M-003902-01-0005). pcDNA3, FLAG-TET2 (#60939) and FLAG-SMAD4 (#80888) were purchased from Addgene.

### Protein Isolation, SDS-Page and Western Blotting

Harvested cell pellets were lysed in Laemmli buffer (50mM Tris-HCL pH6.8, 2% [w/v] SDS, 1X Protease Inhibitor cocktail, 1X Phosphatase inhibitor cocktail) and homogenised using a 23 gauge syringe needle. Extracts were analysed by SDS-PAGE using 4-12% Bis-Tris NuPAGE gels (Invitrogen) and transferred onto PVDF membranes (Millipore). After washing in PBS containing 1% Tween-20 (PBS-T), membranes were blocked in 5% skimmed milk in PBS-T and then incubates with the primary antibody overnight at 4C. Antibodies and dilutions used are Actin (Santa Cruz, 1:5000), AMPK (CST, 1:1000), pAMPK T172 (CST, 1:500), E-Cadherin (CST, 1:1000), GATA6 (Abcam, 1:500), PDX1 (Thermo Fisher, 1:500), SMAD4 (CST, 1:1000), TET2 (Abcam, 1:500). The membranes were incubated with HRP-conjugated secondary antibodies (CST) for 1 h at room temperature and exposed to X-ray film (Kodak) after incubation with Thermo Scientific Pierce ECL or Amersham ECL (GE Healthcare). ImageJ software (NIH) was used for the quantification of the bands. All bands were normalised against the loading controls.

### Dot Blot

Cell pellets were lysed in DNA lysis buffer (100mM Tris-HCL [pH8.5], 5mM EDTA, 0.2% SDS, 100mM NaCL, 0.5mg/ml Proteinase K) and incubated at 55 °C whilst shaking for 72 hours. gDNA was precipitated by ethanol precipitation and DNA pellets were resuspended in nuclease free water. RNAse A digestion was carried out at 37 °C for 2 hours. gDNA was sonicated in a Diagenode biorupter plus for 20 cycles of 30 seconds on, 30 seconds off at high power. DNA was then quantified using a NanoDrop 1000 (Thermo Fisher). For dot blots, DNA was normalized and then denatured by adding 1 volume DNA denaturing buffer (200mM NaOH, 20mM EDTA) and heating at 95 °C for 10 minutes. 2 volumes 20X saline-sodium citrate (SSC) buffer was added and samples were chilled on ice for 5 minutes. Serial dilutions of DNA were directly pipetted onto Amersham Hybond-N+ blotting paper (GE Healthcare) and UV crosslinked at 1200J/m2. Membranes were blocked with 5% milk/PBST and incubated with 1:1000 5’hmC primary antibody (Active Motif, 39769) overnight at 4°C. Membranes were then washed and developed in the same method used for western blot as above. To visualize total DNA membranes were stained with 0.1% Methylene blue.

### qPCR

RNA isolation and cDNA synthesis was performed using Power SYBR Green Cells to-Ct Kit (Thermo Fisher) following the standard protocol. 50,000 cells were lysed in 25uL Lysis solution containing DNase, and 5ul of this lysis used for cDNA synthesis. Target gene expression was assessed using Power SYBR Green PCR Master Mix (Thermo) using primers designed by PrimerBank (https://pga.mgh.harvard.edu/primerbank/. All reactions were performed in duplicate and ran for 45 cycles. Fold change in target mRNA expression was calculated based on the cycle threshold (CT) method after normalization to GAPDH expression. Primers used were GAPDH (fw: GGAGCGAGATCCCTCCAAAAT, rev: GGCTGTTGTCATACTTCTCATGG), SMAD4 (fw: CTCATGTGATCTATGCCCGTC, rev: AGGTGATACAACTCGTTCGTAGT) and GATA6 (fw: CTCAGTTCCTACGCTTCGCAT, rev: GTCGAGGTCAGTGAACAGCA).

### Migration Assays

PANC1 cells were treated daily for 3 days and then switched to medium containing 0.1% serum plus the appropriate treatment and incubated overnight. The following day, cells were harvested, washed in medium containing 0.1% serum and seeded into the upper wells of a CIM-plate 16 (ACEA Biosciences, Inc.) as per the manufacturer’s protocol and at a density of 4 x 10^4^ cells/well. The lower wells contained 10% serum medium. Migration was followed for 30 hr and the time taken to reach peak migration (t1/2) was determined from the first derivative of the migration curve.

